# Gene-methylation interactions: Discovering region-wise DNA methylation levels that modify SNP-associated disease risk

**DOI:** 10.1101/593053

**Authors:** Julia Romanowska, Øystein A. Haaland, Astanand Jugessur, Miriam Gjerdevik, Zongli Xu, Jack Taylor, Allen J. Wilcox, Inge Jonassen, Rolv Terje Lie, Håkon K. Gjessing

## Abstract

The genetic code is tightly linked to epigenetic instructions as to what genes to express, and when and where to express them. The most studied epigenetic mark is DNA methylation at CpG dinucleotides. Today’s technology enables a rapid assessment of DNA sequence and methylation levels at a single-site resolution for hundreds of thousands of sites in the human genome, in thousands of individuals at a time. Recent years have seen a rapid increase in epigenome-wide association studies (EWAS) searching for the causes of risk for genetic diseases that previous genome-wide association studies (GWAS) could not pinpoint. However, those single-omics data analyses led to even more questions and it has become clear that only by integrating data one can get closer to answers. Here, we propose two new methods within genetic association analyses that treat the level of DNA methylation at a given CpG site as environmental exposure. Our analyses search for statistical interactions between a given allele and DNA methylation (*G*×*Me*), and between a parent-of-origin effect and DNA methylation (PoO× Me). The new methods were implemented in the R package Haplin and were tested on a dataset comprising genotype data from mother-father-child triadsm with DNA methylation data from the children only. The phenotype here was orofacial clefts (OFC), a relatively common birth defect in humans, which is known to have a genetic origin and an environmental component possibly mediated by DNA methylation. We found no significant PoO×Me interactions and a few significant G×Me interactions. Our results show that the significance of these interaction effects depends on the genomic region in which the CpGs reside and on the number of strata of methylation level. We demonstrate that, by including the methylation level around the SNP in the analyses, the estimated relative risk of OFC can change significantly. We also discuss the importance of including control data in such analyses. The new methods will be of value for all the researchers who want to explore genome- and epigenome-wide datasets in an integrative manner. Moreover, thanks to the implementation in a popular R package, the methods are easily accessible and enable fast scans of the genome- and epigenome-wide datasets.

## 1 Introduction

The genome includes the code for all the ingredients of the cell, but not every cell needs all the ingredients at all times. The epigenetic mechanisms can modify the level of gene expression without altering the underlying DNA sequence. One of the most widely studied epigenetic marks is DNA methylation^1^, which is also very tightly associated with the DNA code itself.

DNA methylation influences gene expression levels through various mechanisms, not all of which are thoroughly understood^2,3^. In eukaryotes, DNA methylation is a process by which a methyl group is added to a cytosine (C) nucleotide, most often within a CpG (cytosine–phosphate–guanine) dinucleotide motif. The state of methylation is strictly controlled by the activity of several enzymes^4^ and is influenced by the DNA code as well as environmental exposures^3^.

Throughout our history, the human genome has evolved to contain CpG-depleted and CpG-rich regions, the latter often referred to as CpG islands (CGIs)^2,5^. CGIs are mostly unmethylated and located within gene promoter regions where the transcription factors normally bind. Although the methylation of gene promoters can potentially lead to the suppression of gene expression^1^, it has been shown that certain transcription factors actually require their binding sites to be methylated in order to promote binding^6^. In addition to promoters, DNA methylation within gene enhancer regions and gene bodies might also regulate gene expression^1,7–9^. Thus, the function of DNA methylation is highly region-dependent and may vary from gene to gene.

Recent technological breakthroughs in sequencing platforms have enabled the global assessment of DNA methylation levels in thousands of individuals ^10,11^. This has led to a substantial increase in the number of epigenome-wide association studies (EWAS) and attempts to combine EWAS with data from genome-wide association studies (GWAS), resulting in an unprecedented opportunity to improve the mapping of genomic regions that are associated with genetic diseases by incorporating the DNA methylation information. However, EWAS data are relatively new, and although there have been some attempts at standardizing their analysis^12^, there are still a paucity of statistical tools to incorporate DNA methylation data into genetic association analyses^13–15^. Many bioinformatics methods are currently available for analyzing methylation data from genome-wide scans^3,16^, with most of them focusing on identifying the so-called differentially methylated regions (DMRs)^17–20^ and methylation quantitative trait loci (meQTL)^21^. A few studies have used both genomic and methylation data to predict disease prevalence or risk. For instance, Shah et al.^22^ tested several linear models to investigate the extent to which disease prevalence prediction was due to genetic or methylation components alone. Further, White et al. ^23^ performed a multi-step analysis of several data sources (genetic, methylation and transcription) to unravel the association of genes with neurological disorders. In another work, several linear regression models were tested to identify associations between methylation levels at CpG sites and child’s birth weight^24^, and subsequently to test for association between the top CpGs and genotypes. A few studies searched for an association between a specific genotype and the methylation level at a chosen CpG from a neighboring region^25–27^. One study that adopted a truly integrative approach, where all the available genetic and methylation data were taken into account in the analysis, searched for differentially methylated CpG sites and meQTLs associated with breast cancer^28^.

Since DNA methylation may control gene expression by either allowing or preventing specific protein factors to bind to promoters, enhancers or gene bodies, it is natural to ask whether there is an interaction effect between the SNPs that are correlated with a phenotype, and the methylation levels in nearby regions, i.e., whether methylation levels are able to explicitly modify the gene expression relative to disease. In a cohort setting, there were two studies that explored possible G×Me interactions, between genetic variants and DNA methylation^29,30^. The authors analyzed several SNPs and CpGs that lie in a candidate gene for asthma and found one SNP–CpG pair with statistically significant interaction, i.e., the relative risk of asthma associated with this SNP increased with increasing methylation level of this CpG site. However, these studies looked only at a few SNPs within one gene and only at one CpG site, which was chosen based on a significant association with the phenotype.

The concept of gene-methylation interactions becomes particularly relevant in the study of *imprinting effects*. An imprinting effect may be caused by the maternally and paternally derived DNA strands having different methylation profiles, thus potentially leading to differential expression of maternally and paternally inherited SNP alleles. In statistical terms, this would lead to a detectable parent-of-origin (PoO) effect, i.e., a statistical effect where the SNP allele relative risk of disease depends on the parent of origin of the allele.

For a dichotomous phenotype, a commonly employed alternative to the cohort and case-control design is the case–parent triad design, where the cases, as well as their biological parents, have been obtained for genotyping. With the case–parent triad design, PoO effects can be detected and estimated^32^. It is also well known that the design allows standard gene-environment (G×E) interactions to be estimated, in spite of only case individuals having been sampled^33^. Going one step further, it is even possible to evaluate PoO×E-effects, i.e., to discover situations where the parent-of-origin effect is modified by environmental exposure^31,34^.

By considering methylation as “environmental exposure”, the triad design thus allows estimating G×Me and PoO×Me interactions. While G×Me effects would be interpreted as in a cohort or case-control setting, the PoO ×Me interactions offer an intriguing extension. Since a PoO effect can be indicative of imprinting, and since this may happen through differing methylation levels depending on the parent of origin, we expect that methylation levels at nearby CpGs could actively influence the size of the PoO effect. That is, we would expect to detect PoO × Me interaction effects.

In the present study, our objective is to make full use of the case–parent triad design, in which GWAS SNP data are available on mother, father, and child, in combination with EWAS data on the child alone. While parental sources of methylation cannot be identified in this design, it is possible to consider methylation as environmental exposure, and thus analyze G×Me and PoO×Me interactions. Here, we present two new methods that use DNA methylation and genotype data to assess the relative risk (*RR*) associated with a certain SNP or haplotype, and dependent on the methylation level around the SNP. We develop the necessary statistical models to estimate the interactions from triad data and discuss critical assumptions that need to be met to avoid false positives. Moreover, we take into account DNA methylation in regions, not single CpG sites, since it has been shown that CpGs act collectively^8,35^.

While our methods are applicable to any dichotomous phenotype, we focus here on orofacial clefts (OFC) — a relatively common birth defect with a prevalence of approximately 1 out of 800 live births (worldwide average). Many GWAS on clefting have been published and have unraveled a wealth of genes and loci for OFCs; results can be found in a number of reviews and their references^36–39^. However, we know that the environment can influence the risk of OFC^38,40^, and most recently, a Mendelian randomization study provided evidence that it might be DNA methylation that mediates the environmental influence on several genes associated with clefting^41^. Overall, the phenotype is very well suited for testing our new methodology.

## 2 Materials and Methods

### 2.1 Genome-wide data

#### Genotypes

Genotypes on 573 case–parent triads were available from the Norway Facial Clefts Study (https://www.niehs.nih.gov/research/atniehs/labs/epi/studies/ncl/index.cfm). Details of the study paticipants^42^ and information on DNA sequencing and quality control measures used for data cleaning have been published elsewhere^36,43^.

We performed analyses on the genotypes from Norwegian families, focusing on the “cleft lip” subtype of OFC, i.e., isolated cleft lip only (CLO) and cleft lip with or without cleft palate (CL/P). We focused on the following SNPs based on evidence from previous reports showing strong associations between these SNPs and the risk of clefting:

1. rs1443434 in *FOXE1* (Ref.^44^);
2. rs2013162 in *IRF6* (Ref.^36^);
3. rs17820943 close to *MAFB* (Ref.^36^);
4. rs560426 in *ABCA4* (Ref.^36^);
5. rs12543318 and rs987525 on chromosome 8 (Ref.^36,45^).

Moreover, we ran control analyses on non-affected child–mother dyads. DNA methylation data of the children and genetic data of the dyads were available only for the second, fourth and fifth SNP in the list above. The details of data collection and quality control of these samples have been provided in our previous work^46–48^.

#### Methylation data

The Illumina HumanMethylation450 BeadChip (San Diego, CA) was used to assess DNA methylation levels at 485,577 CpG sites, in the blood samples of the children from the triads and dyads mentioned above. Details of raw data processing and quality control are given in the previous work^49^. The methylation level (*β*-value) for each CpG site was calculated as: *β* = *I_M_*/(*I_M_* + *I_U_* + 100), where *I_U_* and *I_M_* are the intensities of the unmethylated or methylated signals, respectively, and the factor 100 is taken to ensure no division by zero. After the quality control, 407, 513 CpGs and 868 samples (456 controls; 105 CLO; 167 CLP and 140 CPO) remained for analysis. Matching of the individuals from the methylation dataset to the genotype dataset yielded 84 and 212 families for the isolated CLO and CL/P phenotypes, respectively. In addition, we were able to match the methylation data to the genotype data from all the 456 control mother-child dyads.

#### Genomic regions

CpG sites located at a maximum distance of 50,000 bp from each SNP were chosen and categorized into specific regions (promoter, enhancer, gene, and, independently, CGI and CpG-poor regions) using the newest information from the ensembl^50^ and UCSC Genome databases^51^. The biomaRt^52^ R package was used to extract information from the ensembl Regulatory Feature database (using GRCh 37), which includes the positions of promoter and enhancer regions in the human genome. Based on this information, we classified a CpG into the category “promoter” when the search returned “promoter” (Sequence Ontology^53^ accession number SO:0000167) or “promoter_flanking_region” (SO:0001952). The category “enhancer” was equal to the “enhancer” description (SO:0000165). Gene regions were derived from the ensembl genome browser (GRCh 37) via biomaRt. CGI regions were downloaded from the UCSC human genome browser (genome version hg19). Because our CpGs and SNPs are based on the human genome hg19/GRCh37 release, we did not use the newest versions of the human genome assembly (hg38/GRCh38) to avoid discrepancies in positional information.

To inspect the methylation data as a whole and check for inconsistencies, we performed two simple analyses: *(i)* statistical summary of methylation levels in the above defined defined regions, and *(ii)* distributions of the methylation levels around the chosen SNPs. More extensive details of these analyses are provided in the Supplementary Material (see Sections S1.1 and S1.2). Overall, the distribution of methylation levels in our data was consistent with other published datasets^54^.

#### Summarizing methylation levels

We averaged the *β*-values from CpG sites located near each SNP, i.e., CpG sites within a gene and CpG sites within a promoter or an enhancer region. Figure 1 presents the distributions of the averaged *β*-values per region for the larger dataset (isolated CL/P). Almost all of the *β*-values had a normal-like shape, except for the gene and promoter regions near rs560426, the promoter region near rs2013162, and the enhancer region near rs987525 and rs2013162. The corresponding distributions for the CLO dataset are similar (see Supplementary Material, Figure S3).

**Figure 1:**
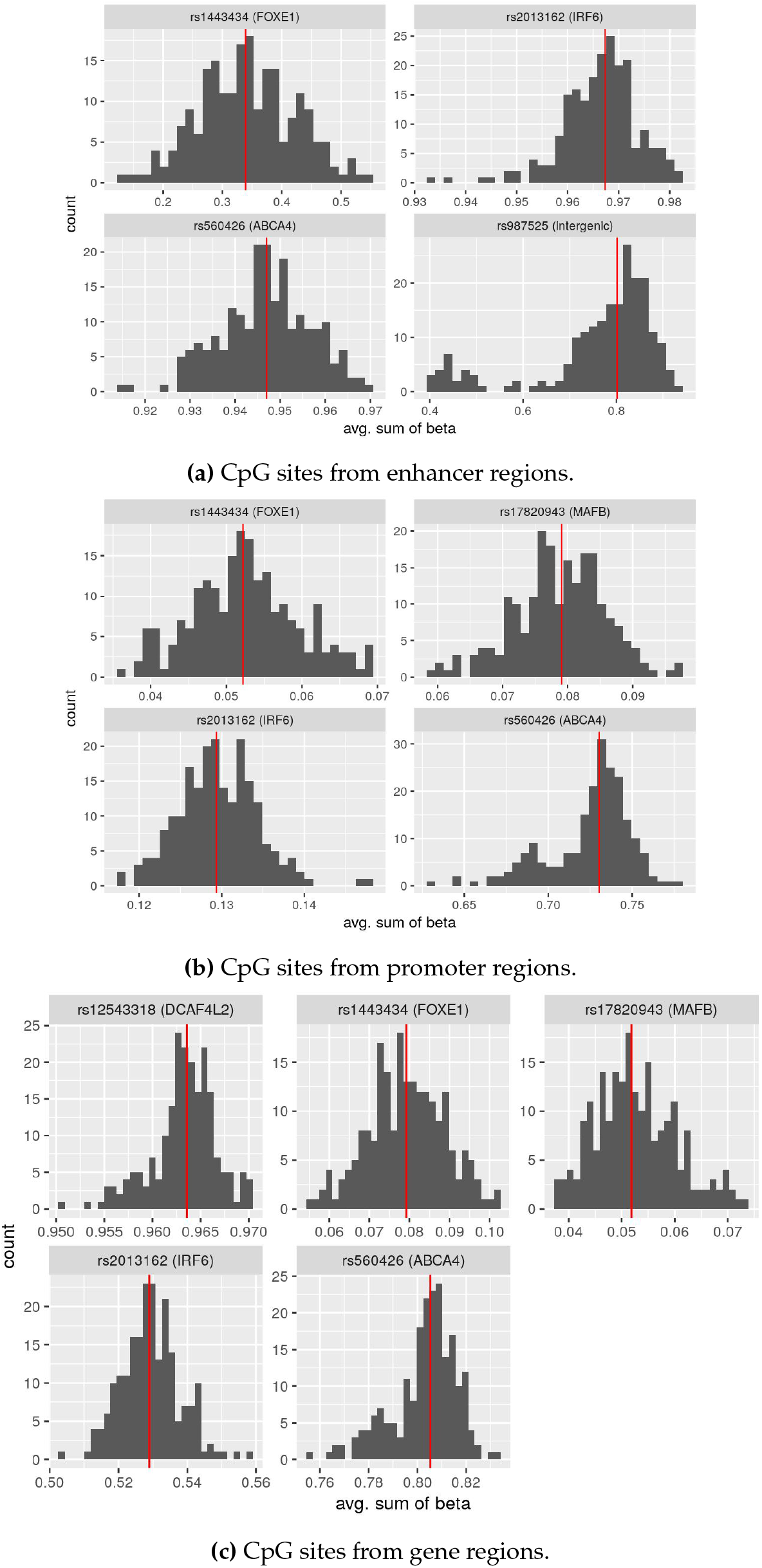
Histograms for the averaged *β* values, CL/P dataset; the red line denotes the median value.

### 2.2 Statistical methods

We applied two different gene–methylation interaction analyses to the case–parent triad data: the G×Me analyses and the PoO×Me analyses. Individuals were stratified into equally-sized groups according to summarized, region-specific *β*-values (low, medium or high). The G×Me interaction analyses compute the relative risk (*RR*) of disease associated with a SNP allele, separately within each stratum of M; a statistically significant change in *RR* across strata indicates a G×Me interaction. The PoO×Me analyses similarly perform PoO analyses within each stratum and then check for a significant change across strata. PoO effects have been described in different ways in the literature — here, we measure it as a ratio of relative risk (*RRR*); if *RR_M_* and *RR_F_* are the relative risks associated with the maternally and paternally derived alleles, respectively, the PoO effect within a stratum is *RRR* = *RR_M_*/*RR_F_*. See Gjerdevik et al.^31^ for further details on the PoO definition. The statistical models are formulated in more detail in the following sections. The strategy for data preparation and analysis is illustrated in Figure 2. As a sensitivity analysis, we ran a four-strata analysis in addition to the three-strata interaction, to see if this influenced the effect detection.

**Figure 2:**
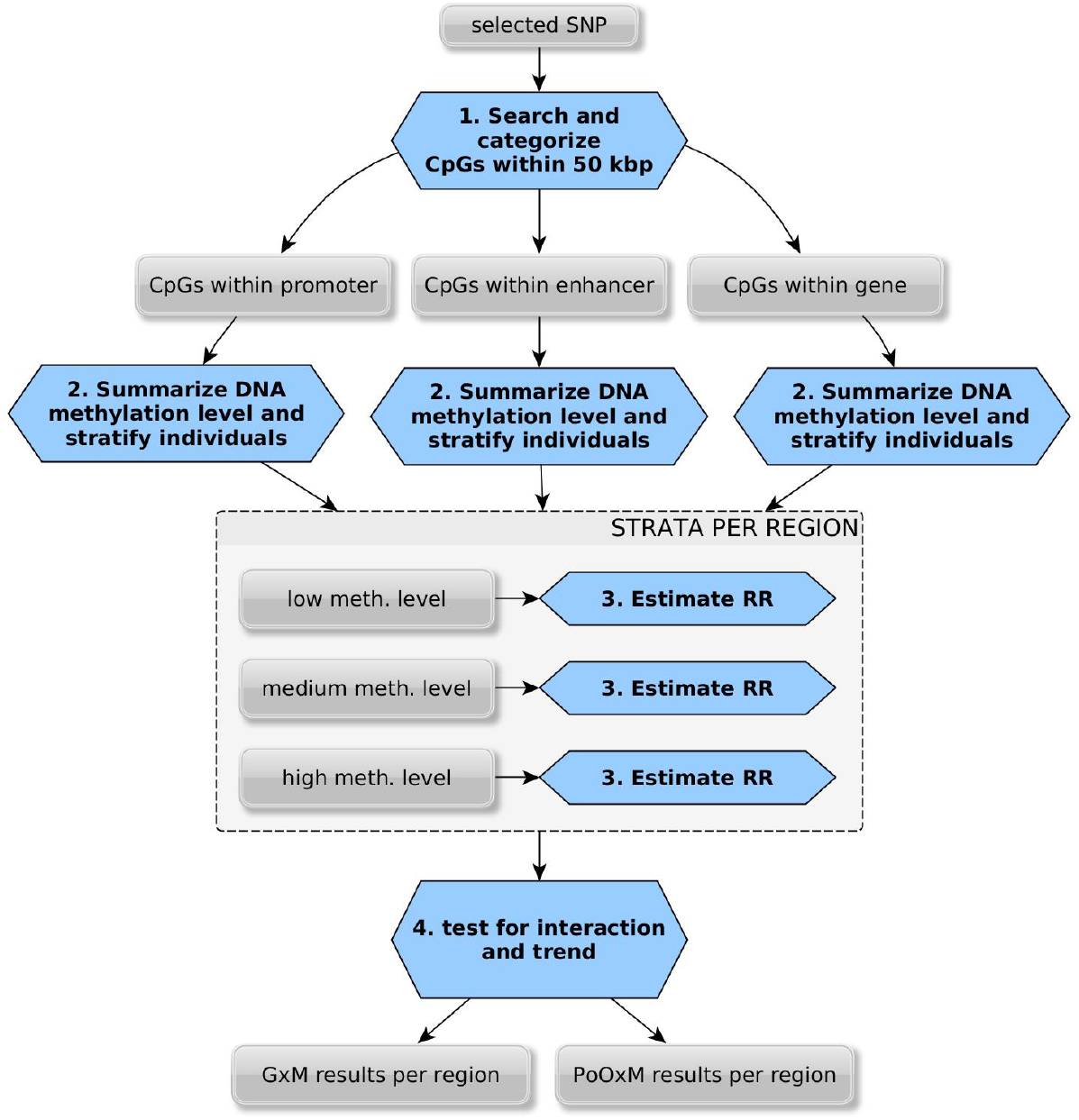
Flowchart of the method for integrating DNA methylation information into genetic association analyses. *RR* = relative risk; G×Me = gene–methylation interaction; PoO×Me = parent-of-origin–methylation interaction.

We also conducted a genome-wide scan on the larger dataset (CL/P) to search for any significant PoO effects. The top 100 results (i.e., the ones with the lowest *p*-values) were then taken as input to additional PoO×Me analyses.

Our Haplin software^31,55^ was used to calculate relative risks with confidence intervals for each of the six SNPs for each phenotype using a log-linear model (`haplinStrat` function), as well as to assess the significance of the interaction, using a Wald test (`gxe` function). The PoO GWAS scan was done using the `haplinSlide` function, with the parameter for ‘window size’ equal to one. While Haplin is available at CRAN, the complete R code used for the analyses is currently available from the authors by request and will be uploaded online.

#### G × Me interactions

Let *M, F*, and *C* denote the genotypes of the mother, the father, and the child, respectively, within a family triad. Here, *M, F*, and *C* will refer to a single SNP, but could also refer to, for instance, haplotypes built from a few SNPs in close linkage disequilibrium. In particular, let *C_ij_* = *A_i_A_j_* denote an ordered child genotype, where allele *A_i_* is inherited from the mother and *A_j_* from the father. *C* denotes the corresponding unordered genotype. Let *D* denote that the child has the disease in question and 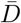 that the child is healthy. Let *Me* denote methylation levels in the child, summarized and grouped as described. We refer to the values when *Me* = *m* as stratum *m*. The penetrance function is *P*(*D*|*C_ij_, Me*), which describes how the probability of disease depends on the child genotype as well as the level of methylation. The penetrance may possibly depend on parent-of-origin since *P*(*D*|*C_ij_, Me*) may differ from *P*(*D*|*C_ji_, Me*). Specifically, we assume that

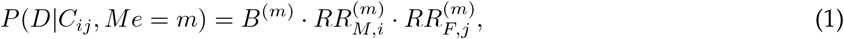

where *B*^(*m*)^ is a baseline risk in stratum *m*, 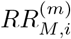 is the relative risk associated with inheriting allele *A_i_* from the mother in stratum *m*, and similarly 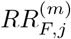 is the relative risk associated with inheriting allele *A_j_* from the father in stratum *m*. We thus assume a multiplicative dose-response model for the allele dose. To make parameters identifiable, we assume 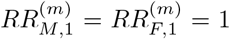 for all *m*, i.e. we choose *A*_1_ as the reference allele in all strata; 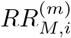 and 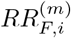 should be estimated from the model when *i* ≠ 1. More details on models that allow deviations from the multiplicative dose-response can be found elsewhere^55,56^.

To look for G×Me interactions, we first assume that 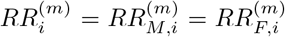 for all *i* and *m*, i.e. that the risk does not depend on the parent of origin. A G× Me interaction effect on the risk of disease would mean that 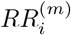 changes over strata of *Me* = *m*. Our data are sampled as case-parent triads and control-parent triads, which means we only observe the distributions *P*(*M, F, C, Me*|*D*) and 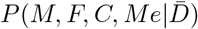, that is, the joint distributions of triad genotypes and methylation strata among the case-parent triads and control-parent triads separately. The number of triads in each category of genotypes and methylation strata are modeled as a log-linear model^55,56^. To assess interactions, the data can be utilized in a number of ways. We start by briefly mentioning standard approaches to estimating G× Me interactions without using parental data.

#### Case and control children

Under a rare disease assumption, 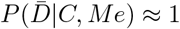. We have

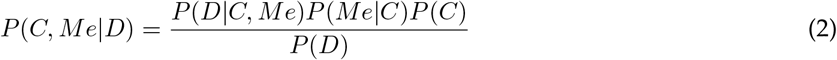

and

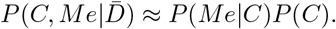

Thus, we can compute the penetrance as the ratio

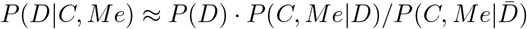

for all combination of *C* and *Me*, which simply corresponds to the standard evaluation of interactions within a logistic regression analysis.

#### Case children only

The relation (2) can by itself be used to assess interaction, under the additional assumption that *P*(*Me*|*C*) = *P*(*Me*), i.e. that genotype and methylation are independent in the background population. If this is the case, we can compute, for instance, the ratio 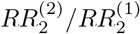 from the corresponding ratios of Equation (2). Note that the relative risks 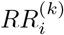 themselves cannot be computed. This interaction computation is the well-known case only-design^33^.

#### Case-parent triads

While the case-control design may have higher power than a case-parent triad design under some circumstances, the case-parent triad design is more resilient to population stratification. In our data, we do not have controls for all SNPs, but case-parent triad data allow interactions to be assessed without controls, and with less restrictive assumptions than the case-only design. Extending (2) to include parents, we can write

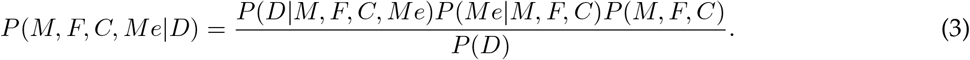

In the standard log-linear model it is typically assumed that the penetrance does not depend on parental genotypes, i.e. *P*(*D*|*M, F, C*) = *P*(*D*|*C*)^55,56^. While this will not be the case if, for instance, maternal genes are involved in disease risk, it is often a tenable assumption. Since *Me* measures methylation in the child, not the parents, it seems reasonable to extend this to the condition

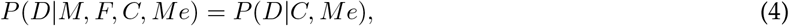

thus ignoring parental genotypes. As in (2), assuming that *P*(*Me*|*M, F, C*) = *P*(*Me*), i.e. that methylation is independent of triad genotypes, the case-triad formula (3) would reduce to the case children formula (2). However, with parents available, we can make a much less stringent assumption. In fact, it is sufficient to assume that

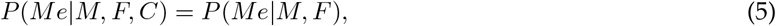

i.e. that methylation in the child may be associated with (parental) genotypes in the population, but that the child genotype does not directly influence methylation. Under the conditions in equations (4) and (5), Equation (3) becomes

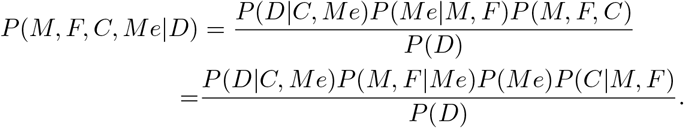

In that case, the term *P*(*D*|*C, Me*)*P*(*M, F*|*Me*)*P*(*C*|*M, F*) corresponds to a stratum-specific log-linear model. *P*(*Me*) and *P*(*D*) are constant within each stratum and are thus irrelevant to the stratum-specific model.

While the case only-design needs the assumption of independence between genes and methylation and is thus prone to population stratification, conditioning on parental genotypes prevents a population association between genes and methylation from distorting the interaction analyses. This is a reasonable assumption made in standard *G*×*E* analyses based on case-parent triads^57^. It should also be noted that the parental genotypes may have a different within-stratum distribution *P*(*M, F*|*Me*) than *P*(*M, F*) in the population at large, precisely if there is, for instance, population stratification that causes a population association between genotypes and methylation. It might also be caused by, for instance, maternal genes *M* that increase the mother’s proneness to smoking, which again influences her child’s methylation patterns. But again, this would not necessarily break condition (5), since maternal smoking habits may only be related to her own genes, and not to which allele is passed on to her child.

However, in our situation, where the “environmental exposure” is methylation (*Me*), it is conceivable that this condition does not hold in situations where the methylation levels at a CpG are directly influenced by a nearby SNP, as is the case for meQTLs.

#### Using the full hybrid design of case-parent and control-parent triads

In cases where control-parent triads are available in addition to case-parent triads, and under the rare disease assumption, we can use the relation

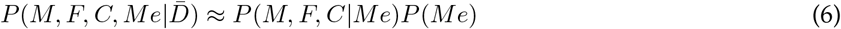

to estimate *P*(*M, F, C*|*Me*) directly among controls, thus eliminating the need for condition (5) altogether. Control triads may thus be used just to check if there is an interaction pattern among controls, which would mean that it is most likely due to an meQTL. Alternatively, the estimate obtained from relation (3) could be adjusted to account for patterns among controls, by including both case and control triads in a hybrid analysis. In the present setting, we use control-parent triads, when available, as independent checks of plausibility.

#### PoO×Me interactions

To estimate parent-of-origin effects, we now assume that *RR_M,i_* is not necessarily equal to *RR_F,i_*, i.e. the effect of allele *A_i_* on offspring risk may depend on which parent it was transmitted from. Let *RRR_i_* = *RR_M,i_*/*RR_F,i_* be the ratio of the relative risks associated with allele *A_i_*. Thus, *RRR_i_* ≠ 1 is indicative of a parent-of-origin effect. Let 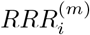 be the value of *RRR_i_* estimated in stratum *Me* = *m*, and *C_ij_* as defined above. As before, we assume *P*(*D*|*M, F, C_ij_, Me*) = *P*(*D*|*C_ij_, Me*). To be able to estimate the ratio 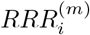 within each stratum, we again need to make an assumption about the term *P*(*Me*|*M, F, C_ij_*), as in relation (5). However, it will now suffice to assume that

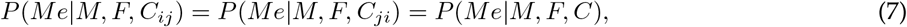

i.e. child genes may influence methylation levels directly, even within parental mating types, but the mechanisms by which the child genotype influences methylation values should not depend on whether specific alleles were inherited from the mother or from the father. Under condition (7),

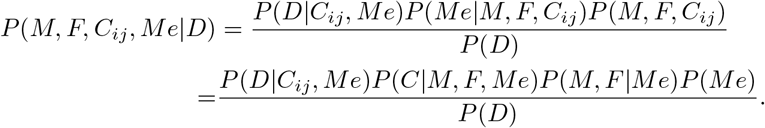

In a log-linear model, we can still obtain estimates of 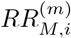 and 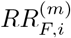 for each stratum *m*. These estimates may be biased since *P*(*C*|*M, F, Me*) may depend on both *Me* = *m* and the (unordered) *C* = *A_i_A_j_*. However, the ratio 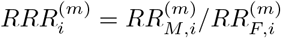 will be unbiased as long as condition (7) is satisfied.

Again, if control trios are available, under the rare disease assumption we can use estimates from the control trios to check the symmetry assumption (7), or in a combined (hybrid) analysis, unbiased estimates of 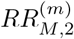 and 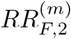 can be obtained.

### 2.3 Presentation of results

To explore these two methods further, we experimented with different setups for the G×Me and PoO×Me analyses. The results are presented in several categories and sub-categories (Table 1). Not all of the regions were included in each of the calculation due to, for example, not having both the methylation and the genotype data, or that none of the identified CpGs were within the specified region. The results are presented in two ways: *(i)* as a quantile–quantile (Q-Q) plot to summarize all the results from different regions, and *(ii)* as a relative-risk plot to show how strong an interaction was for a specific CpG site. We used the R packages ggplot2^58^, gViz [REF], and ggrepel^59^ to create plots. The flowchart (Figure 2) was created using the yEd software (https://www.yworks.com/products/yed).

**Table 1:**
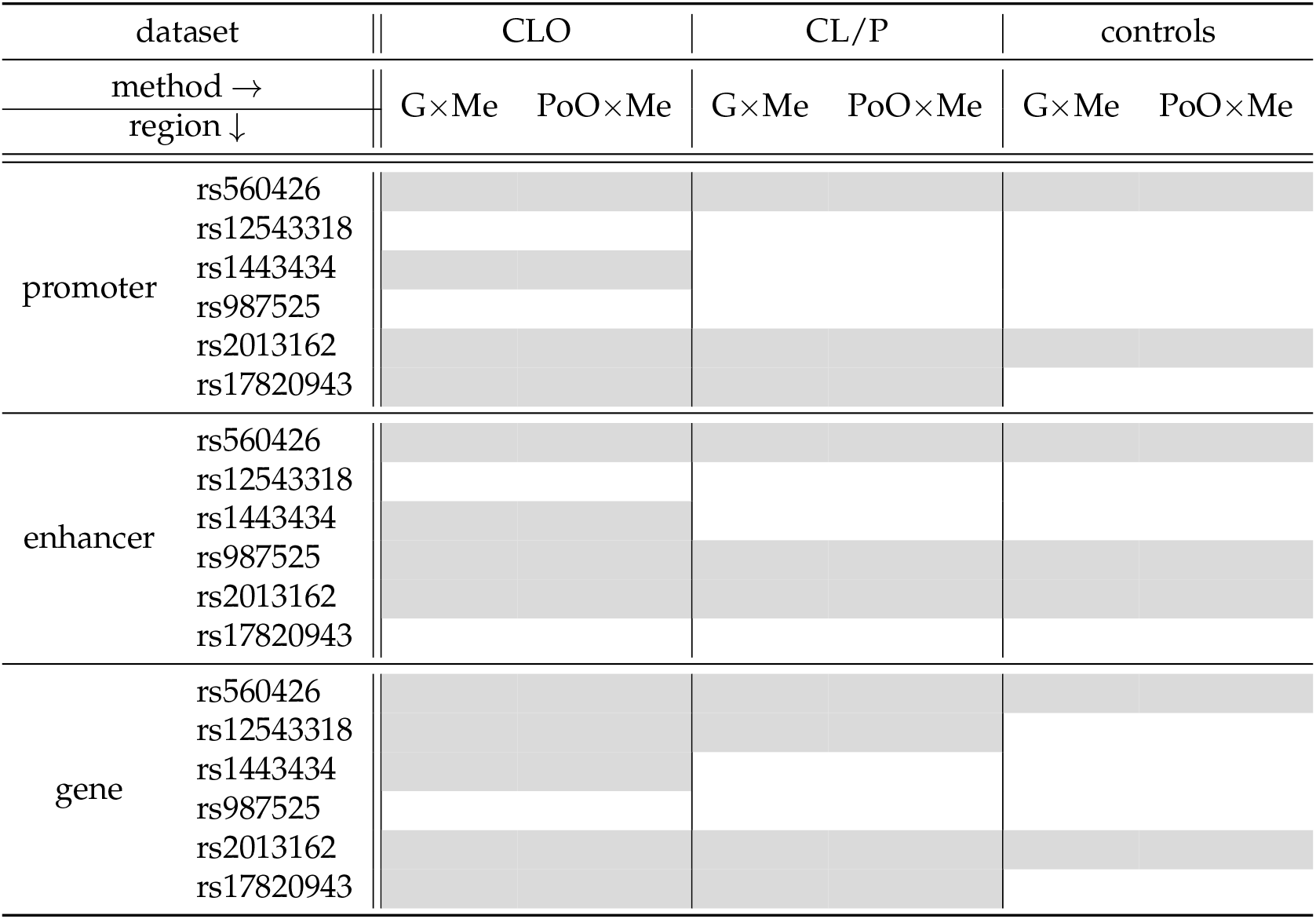
Analyses performed and presented in this article. The grey areas indicate that data were available. We used three genotyped datasets that included individuals with different cleft subtype (CLO or CL/P) and controls, and a DNA methylation dataset, where we classified CpGs into gene, promoter or enhancer classes (see Methods for details).

## Results

We used the averaged *β*-values for all the CpG sites within each defined region (gene, enhancer or promoter) to create different strata and divided the individuals into three equally sized groups of methylation levels: low, medium and high. We ran `haplin` analyses on each stratum separately, and jointly for all the individuals. We calculated the standard relative risk ratio for G×Me and PoO×Me effects. Below, we present the most significant results. All *p*-values are provided in the Supplementary Material, Tables S2–S8.

### 3.1 G×Me analyses

#### 3.1.1 Comparison of quantile–quantile plots

The results of the G×Me analyses are easiest to compare when the Wald interaction test *p*-values are plotted as Q-Q plots (Figure 3). The most significant results in the isolated CL/P dataset were: *(a)* the interaction between rs987525 and the CpGs in the enhancer region nearby; and *(b)* the interaction between rs2013162 in *IRF6* and the CpGs in the gene region. Intriguingly, these interactions were also the first and third most significant in the *control* dataset. We thus consider these results to most likely be false positives. However, the interactions between rs560426 in *ABCA4* and the CpGs in the promoter or surrounding gene regions were significant only in the CL/P and CLO datasets. On the other hand, in the control dataset, a significant interaction was found between rs560426 in *ABCA4* and the surrounding CpGs in the enhancer region.

**Figure 3:**
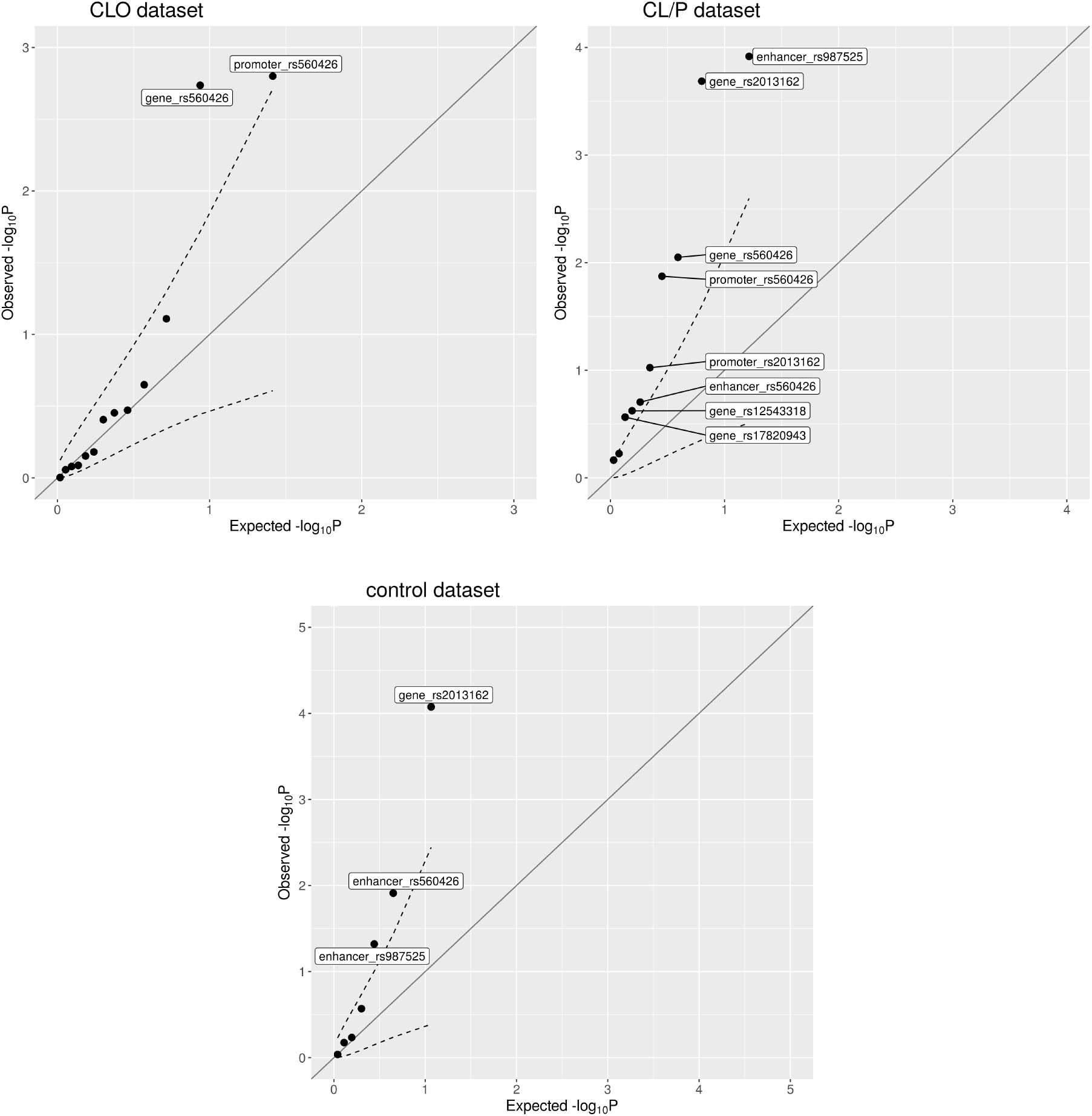
Quantile-quantile plots of the interaction *p*-values from the G×Me analyses; the dashed lines represent the 95% confidence interval.

#### 3.1.2 Detailed results for the top hits from G×Me

##### SNP rs2013162 in *IRF6* showed a significant interaction with methylation levels in the gene regions in two datasets

The plot depicting how the relative risk (*RR*) depends on the methylation level (Figure 4) shows a high degree of consistency across the two datasets in terms of methylation level, the change in *RR*, and importantly, the reversal of the *RRR* for a single-dose versus a double-dose of the minor allele at the given SNP. The fact that this trend is also seen in the controls is a strong indication that the interaction may be a false positive. Moreover, this interaction remained significant when we performed analyses on four instead of only three strata (*p*-value for trend was 5.8 · 10^−4^ for the CL/P dataset, and 7.7 · 10^−6^ for the control dataset; see Supplementary Material, Figure S6).

**Figure 4:**
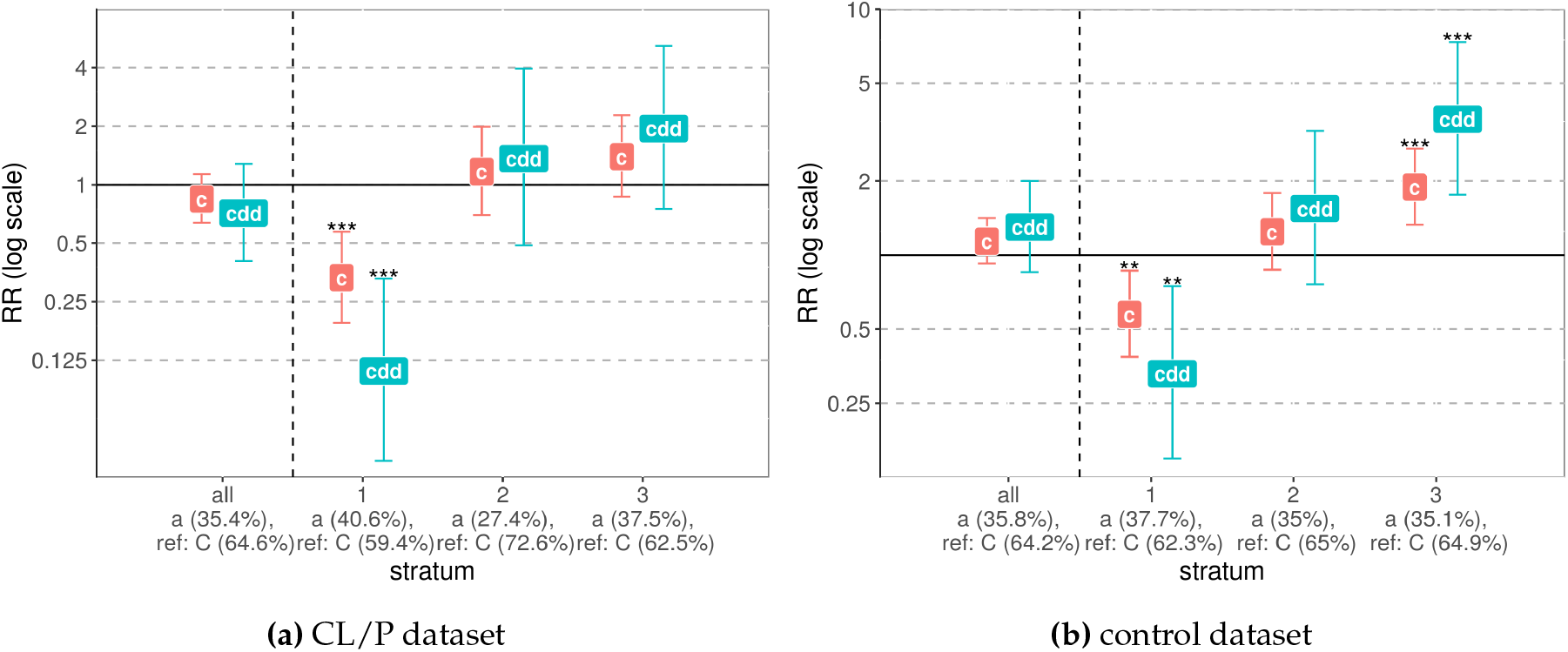
Interaction between a SNP and methylation (G×Me) of the CpGs from the gene region around SNP rs2013162 in *IRF6*. The x-axis groups the results into “all” (without stratification of the dataset) and the results for each stratum: “1” denoting low methylation level, “2” — medium, and “3” — high methylation level. The y-axis shows the relative risk (*RR*) on a log scale, with “c” denoting the child effect when only one minor allele is inherited, and “cdd” denoting the child effect of a double-dose allele inheritance.

Similarly, the interaction between rs987525 and methylation of the surrounding CpGs in the enhancer region was significant in the CL/P and control analyses (see Supplementary Material, Figures S4 and S5b). However, the significance of this interaction in the controls was lost when we increased the number of strata from three to four (Fig. S6).

##### The G×Me analysis of the CLO dataset pointed to interaction with rs560426 in *ABCA4*

The methylation level around this SNP for the CpG sites within both the gene and promoter regions showed a significant interaction with the genotype (Figure 5). Although the overall effect of having a G-allele instead of an A-allele at rs560426 increased the risk of CLO, low methylation level in either gene or promoter regions (stratum 1) decreased the risk. By contrast, among the individuals with high methylation level (stratum 3) in the mentioned regions, the calculated risk was higher than when methylation was not considered at all. Even though the confidence intervals for *RR*s in each stratum were wide, the Wald test for trend indicated that the effect was significant (*p*-value of 8.8 · 10^−5^ and 6.1 · 10^−5^ for the promoter and gene regions, respectively). Moreover, running the same analysis on four strata of methylation level showed that the effect was still significant for these regions (*p*-values for the trend tests were 5.8 · 10^−5^ and 7.4 · 10^−5^ for the promoter and gene regions, respectively, Fig. S6).

**Figure 5:**
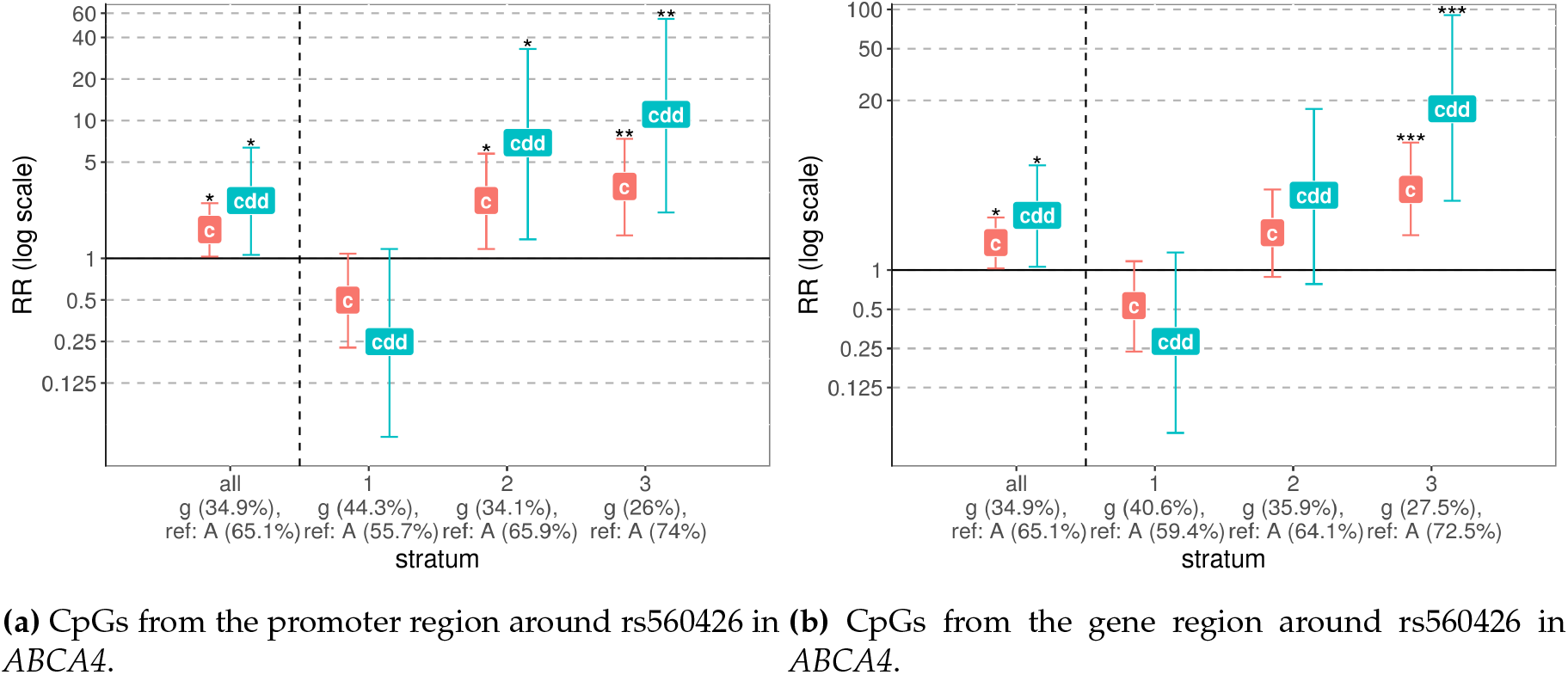
The most significant results of the G×Me analysis of the isolated CLO phenotype. The x-axis groups the results into “all” (without stratification of the dataset) and the results for each stratum: “1” denoting low methylation level, “2” — medium, and “3” — high methylation level. The y-axis shows the relative risk (*RR*) on a log scale, with “c” denoting the child effect when only one minor allele is inherited, and “cdd” denoting the child effect of a double-dose allele inheritance.

#### 3.1.3 Random-SNP analysis

As another test to verify that our method produced reliable results, we ran the G×Me analysis on 20 randomly chosen SNPs (Supplementary Material, Table S5), using the isolated CL/P dataset. As expected, these analyses did not produce any significant *p*-values (Figure 6).

**Figure 6:**
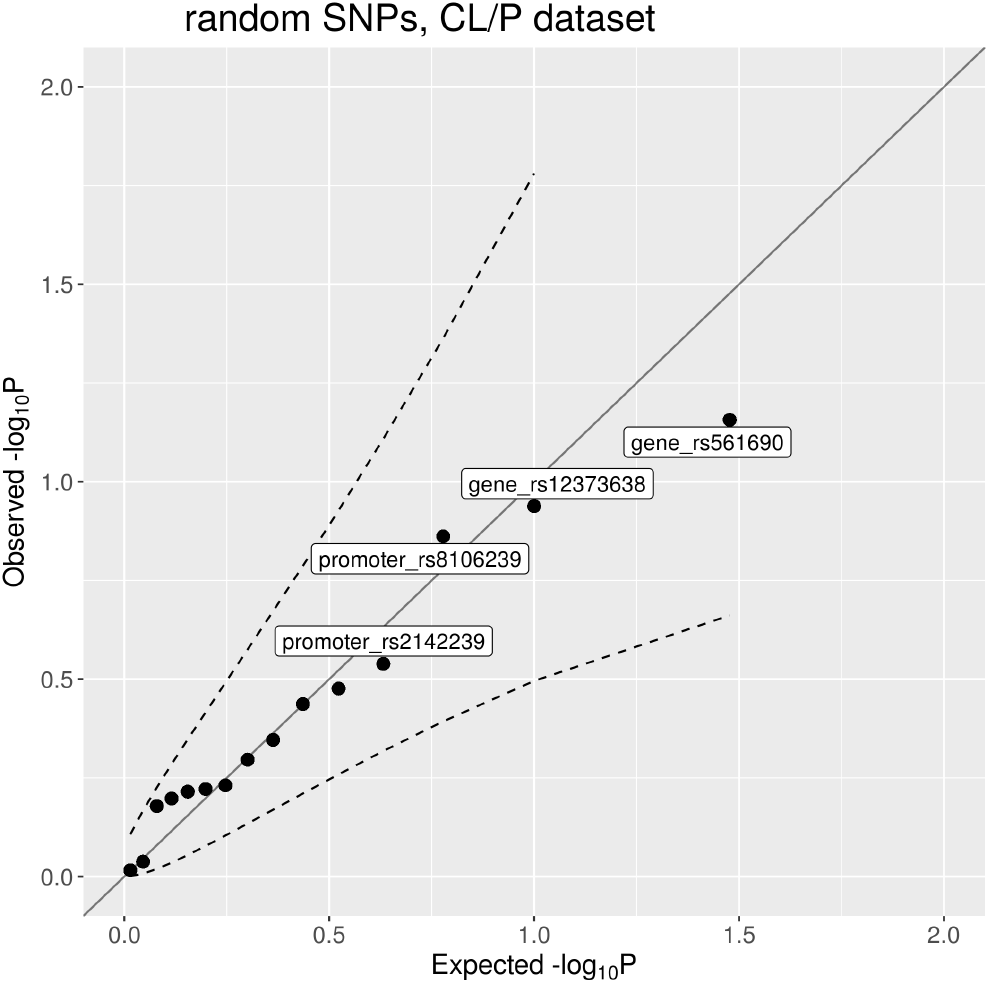
Quantile-quantile plot of the G×Me results from the analysis of 20 randomly chosen SNPs in the isolated CL/P dataset.

### 3.2 PoO × Me analyses

#### Methylation levels did not interact with alleles showing significant PoO effect

There were no significant hits in the PoO×Me analyses (Fig. 7). To obtain further candidates for PoO×Me interactions, we conducted a full PoO GWAS scan of the entire genetic dataset (432 087 SNPs, see Supplementary Material, Table S9). The top 100 hits from the scan had PoO *p*-values in the range (1.8 · 10^−5^; 3.6 · 10^−4^). We then performed subsequent PoO×Me analyses on these top 100 hits. As seen in Figure 8, a number of SNPs exhibited *p*-values more significant than expected by chance; in particular, rs1441744 appears above the QQ-plot 95% pointwise confidence interval for both promotor and gene regions. However, in light of the modest *p*-values and lack of control data for these SNPs, these findings should only be seen as candidates for a possible follow-up.

**Figure 7:**
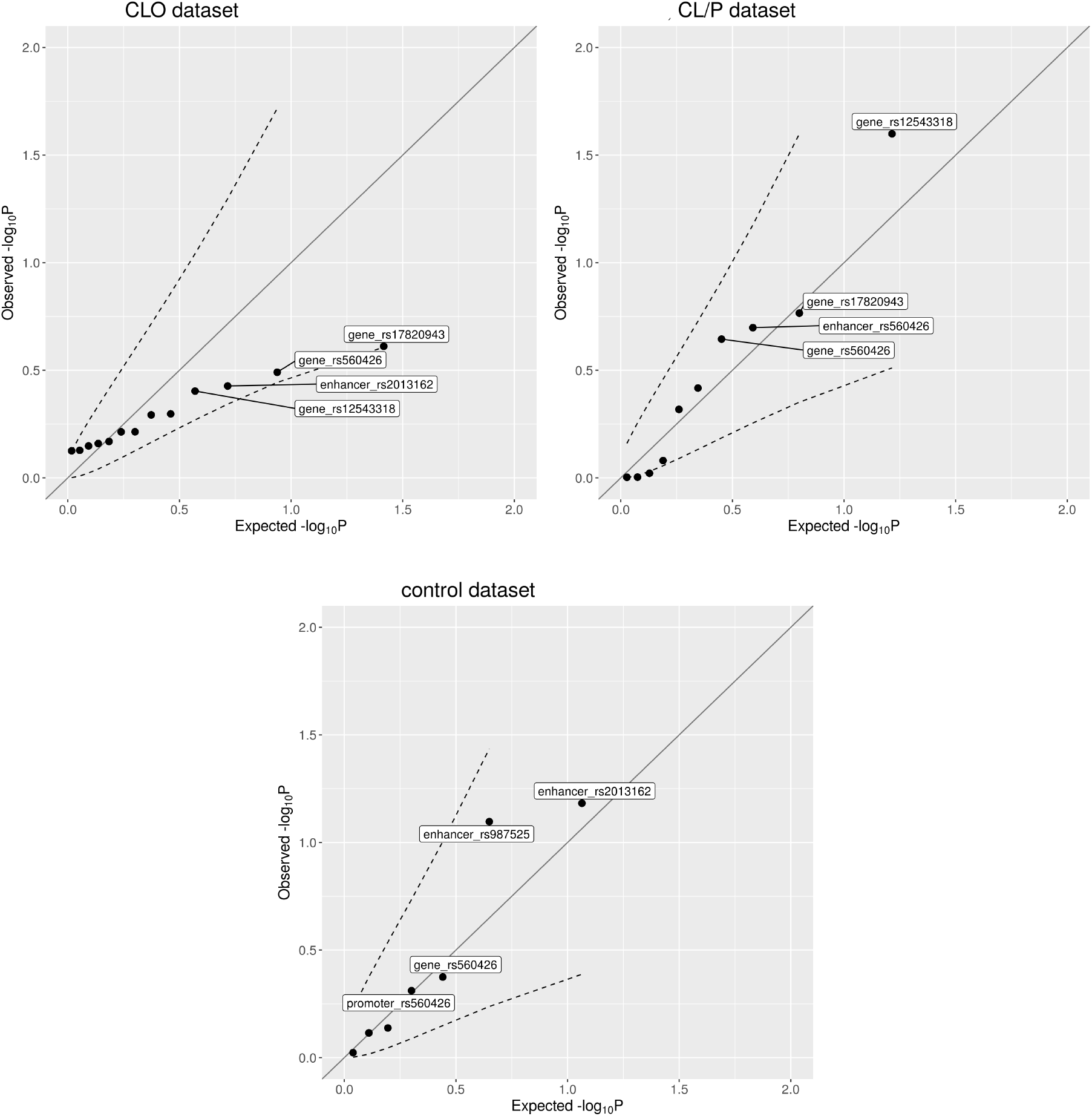
Quantile-quantile plots of the interaction *p*-values from the PoO×Me analyses; the dashed lines represent the 95% confidence interval.

**Figure 8:**
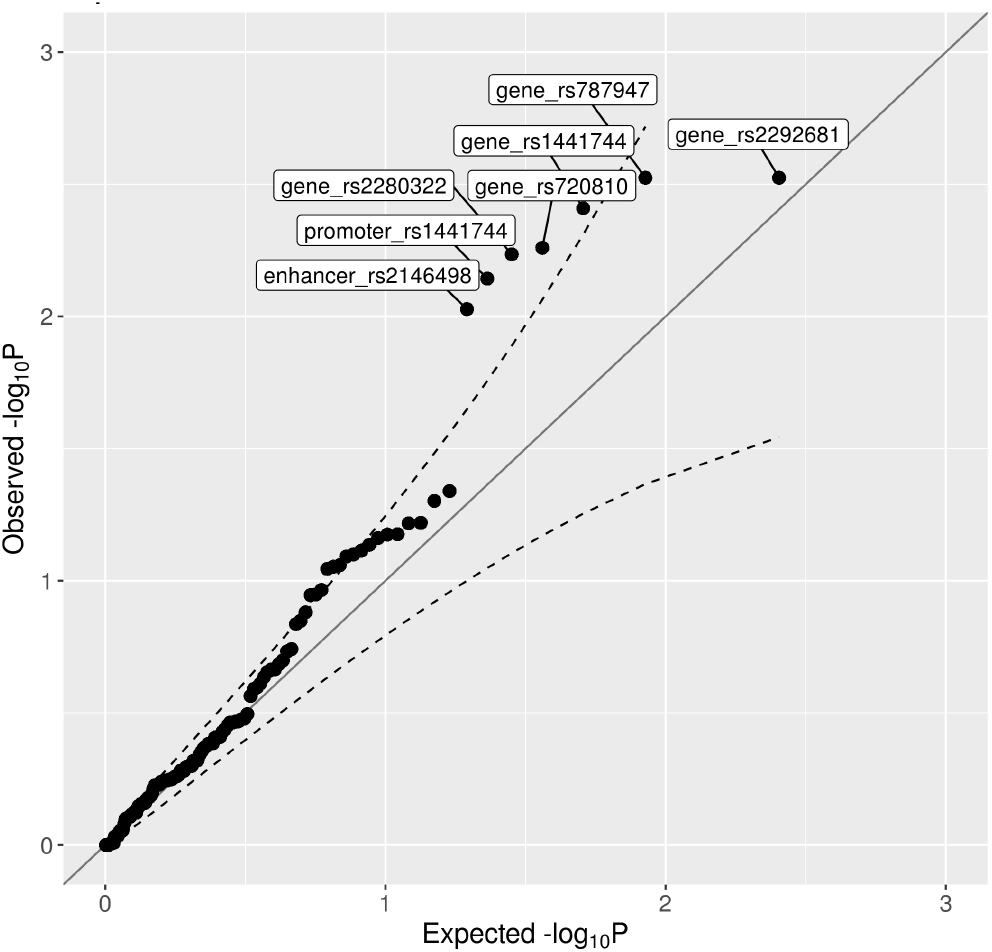
Quantile-quantile plot of the *p*-values from the PoO × Me analysis using the top 100 SNPs with the lowest *p*-values from the GWAS scan for PoO effects; the dashed lines represent the 95% confidence interval.

## 4 Discussion

We presented two methods that incorporate DNA methylation data into genetic association analyses, which we call G×Me and PoO×Me. Although G×Me interactions have been considered previously^29^, we used it here in a much more extended setup. We also included a novel PoO ×Me method that does not have the disadvantages of G× Me. We applied the two methods on the genotypes and methylation data of individuals with cleft lip, as well as of control individuals. We focused our analyses on a set of SNPs that had previously shown significant associations with the risk of clefting. Our results showed that the methods can detect G× Me interactions, but with a possibility of obtaining false positive results. Here, we delve deeper into these issues as well as discuss technical choices regarding the design of our methods.

### Summarizing the methylation patterns in regions

We chose to use methylation level of CpG regions, not only single sites, and used the most straightforward method to average *β*-values from all the CpG sites within a region. This single value from each region was then used to divide the individuals into equally-sized groups, from the lowest to the highest average methylation level. Although each single CpG site can have a unique methylation status, it has been shown that the methylation levels of nearby CpGs is correlated, especially within CGIs^2^. Averaging the methylation level over a region was used e.g., in the search for DMRs^60^, when imputing the missing values^61^, or when defining a methylation score for a region^62^. If there is one CpG site with a much wider range of methylation than the other sites within the region, it will be this specific site that drives the average value. These features of average methylation level make it a good starting point for analyses presented here. We plan to perform further checks of different parameter choices and assumptions of our methods, including how the results are influenced by the choice of methylation summarizing method.

This strategy has one important caveat. For instance, if there were only two CpGs within a given region, and for two persons the *β*-values at these CpG sites were 5 and −5, and −5 and 5, respectively, these two individuals would still be placed in the same methylation category, since the sum is 0 in both cases. However, this effect was not observed in our dataset; overall, the range of *β*-values per CpG site was narrow (Figure S2). The only exception was cg04350215 near rs560426, where the *β*-values ranged from 0 to around 1. Because cg04350215 was the only exception, it is likely that this CpG site dictated the stratum membership for this SNP.

### How to choose the CpGs for the G×Me analysis

Our choice of CpGs is guided by the need to explain the results in a biologically meaningful context. DNA methylation controls gene expression by either allowing or preventing specific protein factors to bind to promoters, enhancers or gene bodies^1,2,9,63^. Therefore, we selected these three regions for our analyses. Importantly, Anastasiadi *et al*. ^8^ were looking at changes in gene expression associated with changes in DNA methylation distribution within promoters, gene bodies, and gene body sub-regions. Their results showed that in the cases where the association was significant, one could basically use mean or median value instead of looking a the distribution. A gene body may encompass a large region, however, it has been shown that those may hide so-called cryptic promoters (see e.g., review^4^), which can regulate levels of other genes when their methylation status changes. Therefore, taking into account all CpGs available within the gene region might increase our statistical power.

We chose 50 kbp as the maximum distance from the SNP for defining the “nearby CpG sites” to incorporate the promoter region and the gene region in the search. The enhancers are known to act also at larger distances, but still the majority of enhancer–gene pairs are within 50 kbp^64,65^. We plan to perform a larger sensitivity analysis of those parameters; however, this is not the topic of this paper.

### The missing data

One limitation is that our analyses are based on data from a microarray chip, which means that we do not have data on all CpG sites within each of the regions considered. However, recent extensive genome-wide analyses of DNA methylation suggest that the chip-based methods provide almost as much information as sequencing techniques^66^.

### Robustness of the approach

We verified the robustness of our method and results by conducting additional analyses: *(i)* by dividing the individuals into four strata of methylation levels instead of just three, and *(ii)* by running the G× Me method on 20 randomly chosen SNPs. The analyses in *(i)* helped to identify the most significant results. The significance of the interaction remain when the number of strata changes. However, the results from three equally sized strata are usually easier to interpret. Moreover, the number of individuals in each stratum should be large enough that the asymptotic properties underlying the Wald tests and likelihood analyses in the log-linear models would be expected to hold true^67^.

On the other hand, the analyses in *(ii)* showed that there is a very low probability of getting false positive results just by chance, since we did not find any significant interactions when randomly picking SNPs from our dataset. However, there might still be false positive results due to specific unmet underlying assumptions, as discussed in the next paragraph.

### G× Me results

As discussed in the Methods (Subsection 2.2), applying the G× E principles to DNA methylation data in a family scenario can lead to false positive results due to a possibly violated assumption of the independence between methylation levels and alleles, even conditional on parental genotypes. Therefore, we sought to apply the same procedure on the control dataset. The results of G× Me on controls gave significant interactions for the rs2013162 SNP in *IRF6* and methylation within both gene and promoter regions around this SNP. This underlines the importance of the use of controls when checking for G×Me interactions. However, the reason for these two interactions within *IRF6* gene being significant in both controls and cases are unknown. It might be that this interaction is not specific to cleft lip since the methylation data we have is measured in cord blood, not in the tissue of interest. Nevertheless, the overall correlation between DNA methylation levels in blood and the lip/palate tissues were found to be high^68^.

Yet another reason for significant interactions of alleles and methylation within the *IRF6* in both cases and controls might be the role of the gene itself. In fact, *IRF6* is a probable DNA-binding transcriptional activator (UniProt ID: O14896, https://www.uniprot.org/uniprot/O14896).

### PoO×Me results

The PoO×Me method presented here has a milder requirement on the independence between methylation levels and genotype. Furthermore, it makes full use of the advantages of the triad design. Therefore, it is well suited for the type of data available. However, since DNA methylation levels represent an average from several cells and, importantly, an average from the two DNA strands, some issues may remain. One is whether we are able to correctly capture the interaction between PoO and the *averaged* methylation level; that is, how sensitive is the PoO effect in relation to the methylation? A recently published study reported a detailed DNA methylome dynamics in the early embryonic development^69^. The authors showed that while in the early development, the paternal DNA methylation pattern may be very different from the maternal one, they found only a few genes whose expression patterns would match the DNA methylation differences. It is likely that DNA methylation alone cannot induce significant changes in expression; however, it is associated with gene expression variation^70^.

Our approach to studying PoO×Me effects is closely related to the contribution of methylation to imprinting. An imprinted locus can be seen as a locus where methylation levels in the child may depend on the parent of origin of the DNA strand. This may potentially lead to an up or down-regulation of the expression of alleles at that strand in a parent-of-origin-specific manner. There have been many approaches to capturing all genes that exhibit imprinting^71^. To identify more loci that are potentially involved in imprinting, a recent study analyzed combinations of DNA methylation and genotypes in the child of mother–child dyads^72^. First, the authors use maternal genotypes to establish, whenever possible, the parent-of-origin status of SNP alleles in the child DNA. Second, they search for SNPs that is associated with methylation status of nearby CpG sites in a parent-of-origin-specific manner. In our notation, this corresponds to finding loci where

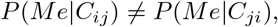

that is, where the distribution of methylation values at the relevant CpG depends not only on the SNP alleles themselves but on their parent of origin. Interestingly, this is closely related to our assumption (7), which we use to exclude possible false positives by checking the condition in control families. While the approach of^72^ is not related to a concrete phenotype, our analyses are specific to a given phenotype; we look for parent-of-origin-specific *Me, C* correlations that are present in case children but not in control children.

### Conclusion

This study implemented two new strategies of searching for interactions between DNA methylation levels and genotype (G×Me) or parent-of-origin effects (PoO×Me). In addition, we show how to use region-wise methylation levels, based on biologically relevant genomic regions: promoter, gene body, and enhancer. The inclusion of these methods in our Haplin R package enables ease of use and adaptation, as the code is open-source and free. We have tested the methods on a modestly-sized group of individuals; however, they can equally well be applied to large sample sizes. We performed a number of sensitivity analyses to ensure the robustness of the approach. While the triad design allows all of the necessary analyses to be performed, we note that independent control triads are required to fully check for – or correct – false positive results.

## Software

Haplin is implemented as a standard package in the statistical software R^73^ and can be installed from the official R package archive, CRAN (https://cran.r-project.org). More details about Haplin can be found on our website, https://people.uib.no/gjessing/genetics/software/haplin.

## Acknowledgements

Support for this work was provided by the Bergen Medical Research Foundation (BMFS) (Grant 807191), and by the Research Council of Norway (RCN) through Biobank Norway (Grant 245464/F50) and the Centres of Excellence funding scheme (Grant 262700). The funding bodies played no role in the design of the study, analysis or interpretation of data, nor in writing the manuscript.

## Supplementary Information

### S1 Methylation distribution

#### S1.1 Methylation levels in promoter, enhancer and CGI regions

We summarized the methylation levels in the following regions: *(i)* CpG islands (CGIs) and *(ii)* CG-poor regions, *(iii)* promoter and *(iv)* enhancer regions. Although the median methylation level was lower in CGIs than in CG-poor regions (Figure S1a), the difference was not significant (the beta values in both these groups spanned the entire range). The promoter regions in our dataset were mostly unmethylated and differed significantly from the highly-methylated non-promoter regions (Figure S1b). Finally, while the enhancer regions were almost all highly methylated, the methylation level of the non-enhancer CpGs spanned the entire range (Figure S1c).

**Figure S1:**
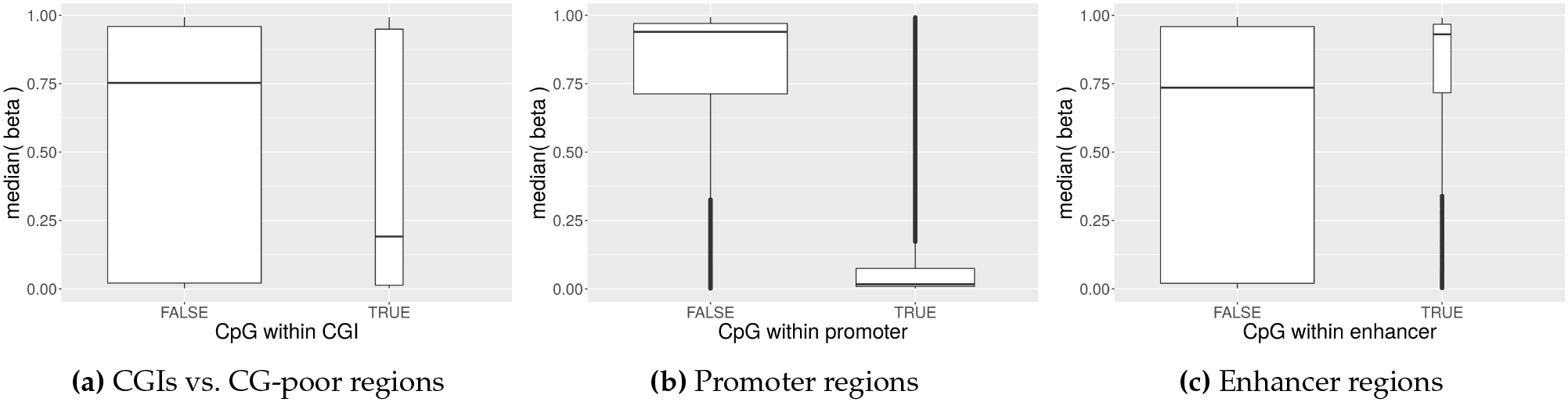
Comparison of median *β*-values from our dataset for specific regions. The width of each box is proportional to the number of observations in the group.

The naturally CG-rich CGIs are expected to be hypomethylated. In contrast, CG-poor regions are not expected to have a specific methylation state, and can even be methylated prior to the binding of transcription factors^1,2^. Promoter regions are expected to be unmethylated in genes that are being transcribed. This differs from enhancer regions, where the methylation was shown to be correlated with increased gene transcription^1,2^. In our dataset, the general methylation patterns of the mentioned regions are consistent with these patterns.

#### S1.2 Methylation levels around the chosen SNPs

We focused our analyses on several SNPs and chose CpGs in their vicinity. First, we analyzed the overall methylation patterns in these regions (Figures S2). Since the data we had came from a selected set of CpG sites on a chip, each of the mentioned categories have a distinct amount of CpG sites by design, with the CpG sites within genes outnumbering those in the other categories (not including the CpGs that do not belong to any of those categories).

Generally, the range of methylation level for each CpG site is quite narrow, with several interesting exceptions, e.g., cg04350215 near rs560426 or cg26035071 near rs2013162 (Figures S2). If we focus on all the CpG sites with an annotated regulatory feature, we can see that their methylation levels span overall a broad range of the *β* values, apart from the data near *DCAF4L2*. These CpG sites belong to the gene region and have a very similar, high methylation level in our data.

**Figure S2:**
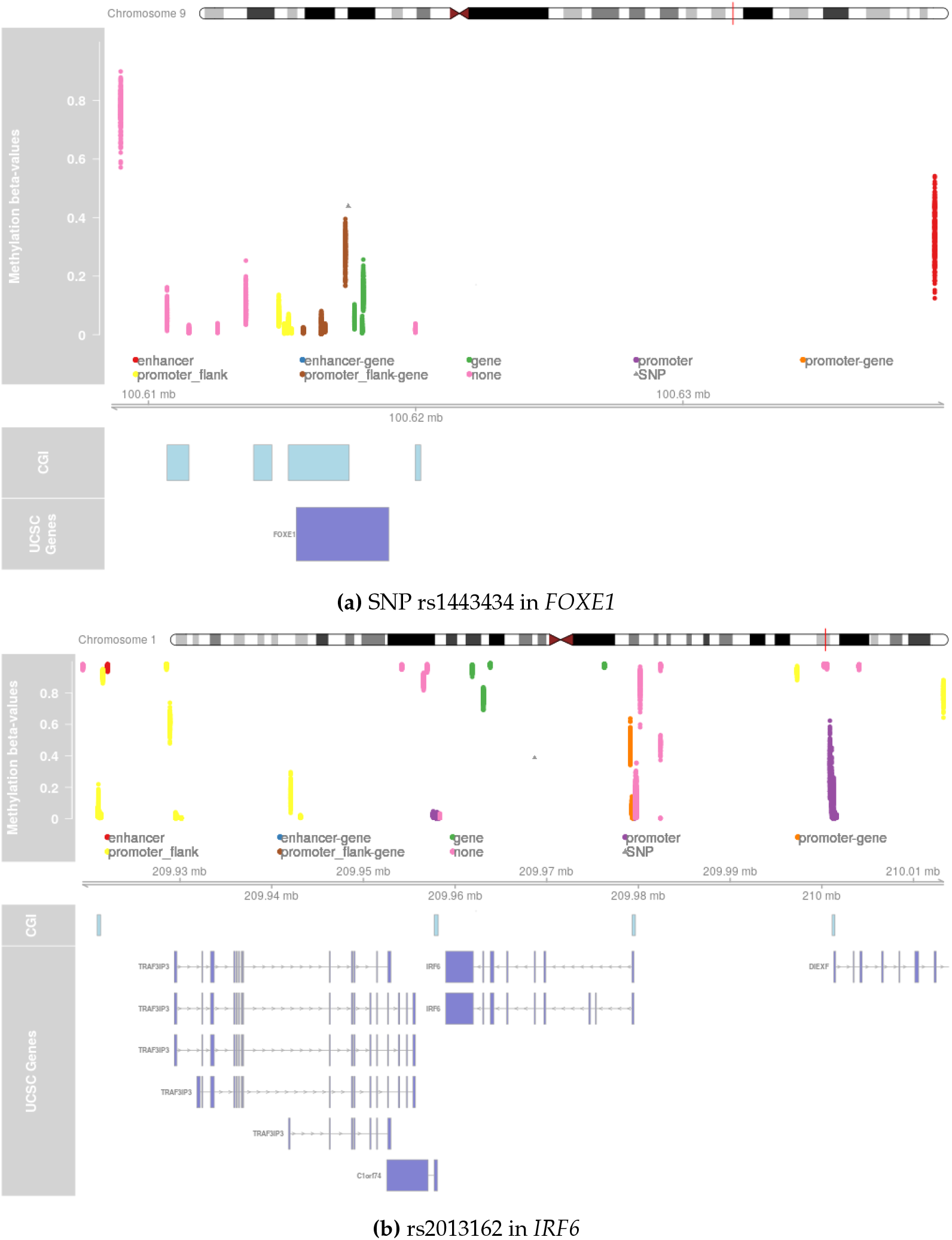
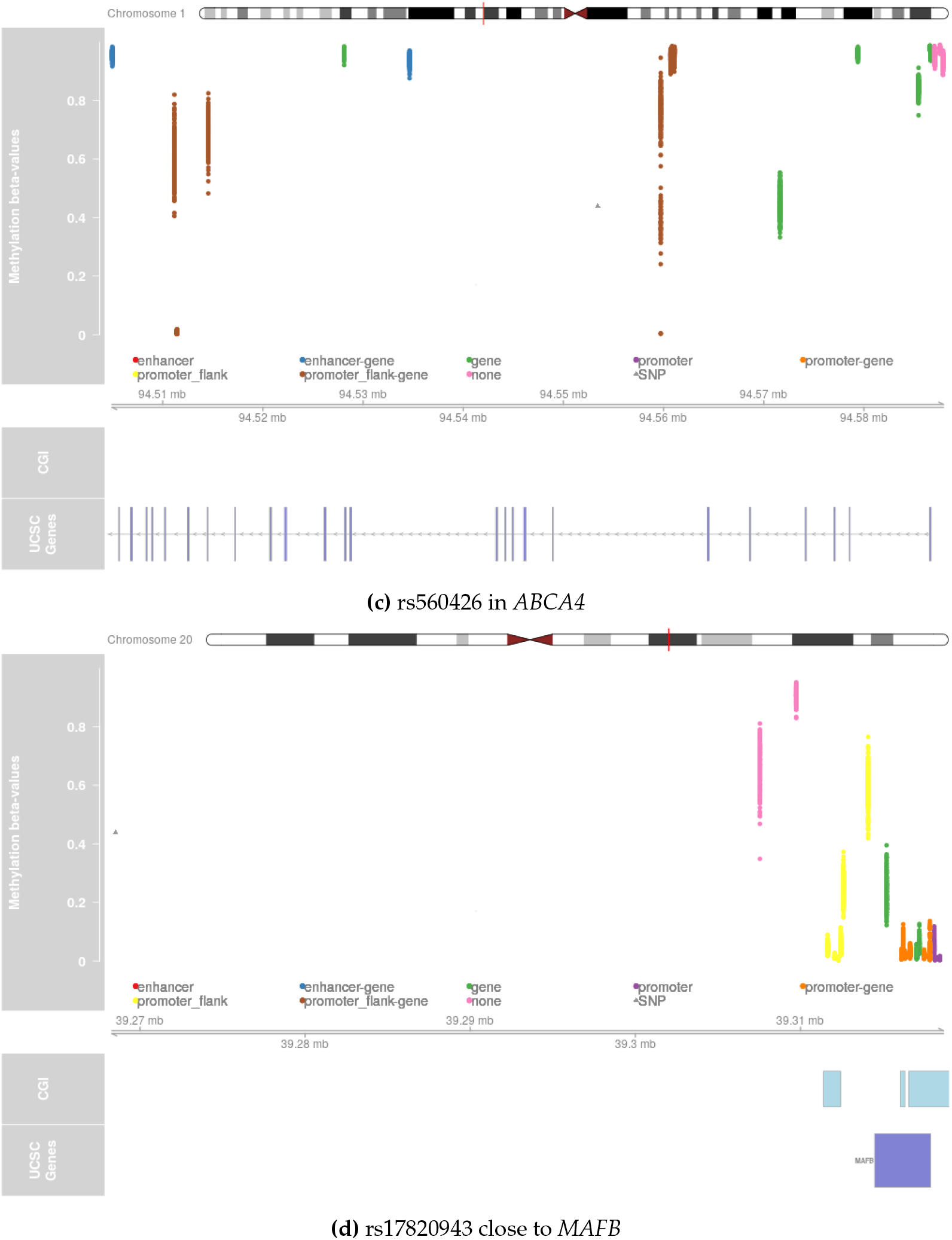
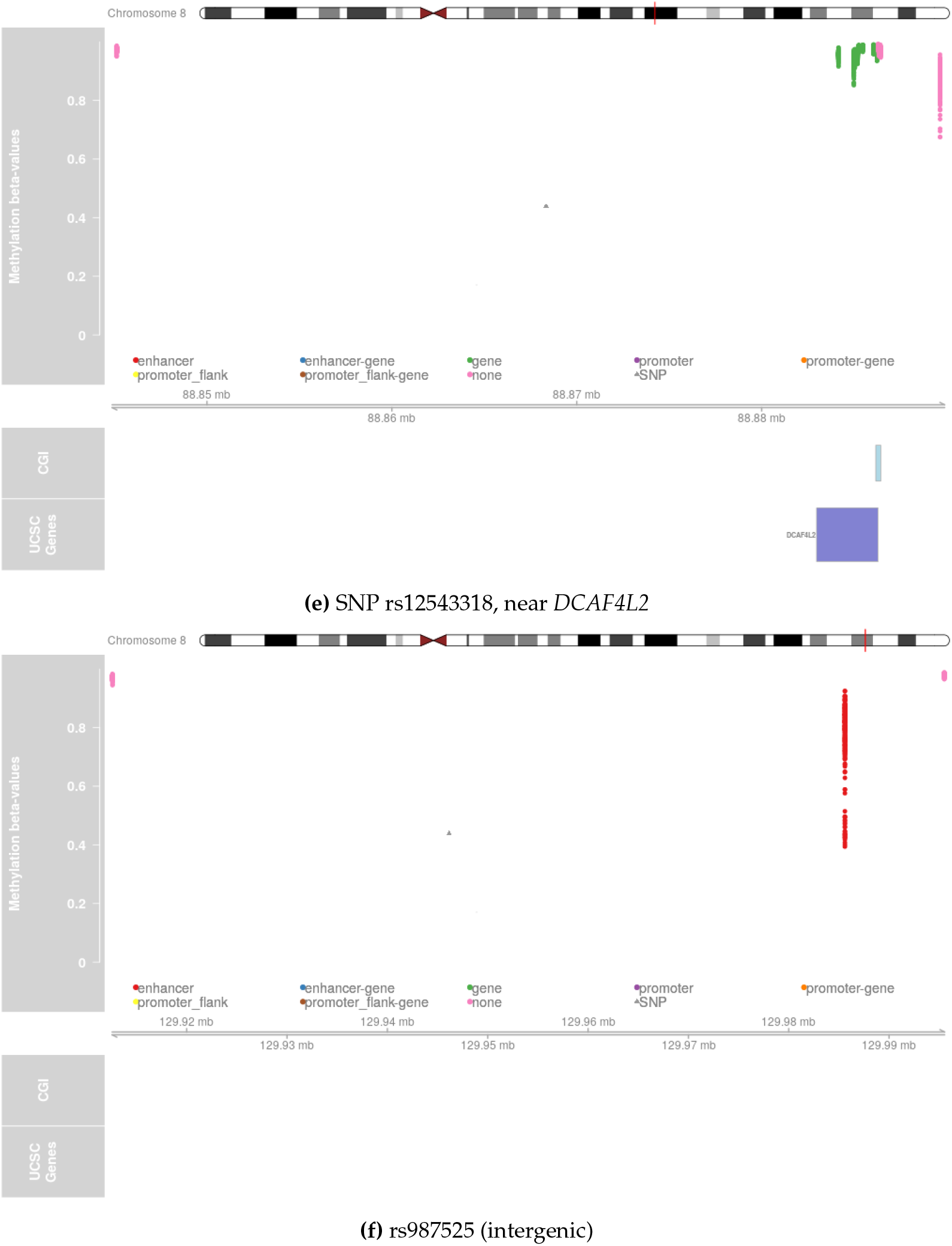
DNA methylation *β*-values for the individuals in the CL/P case-dataset, in the 50 kbp regions around the chosen SNPs. The chromosome ideogram on the top shows the position of the SNP as a red line. Below the chromosome are the *β*-values from all the available individuals; the points are colored according to their regulatory role and if a CpG site was found to be simultaneously in a gene region and in a regulatory region, it was marked as “promoter-gene”, “promoter_flank-gene” or “enhancer-gene”. The axes below the legend shows the positions of each data point, in “mb” (mega base pairs). The two lowest tracks show, respectively, the positions of CpG islands (CGI), if any, and transcripts (UCSC Genes), if any.

#### S1.3 Distributions of the *β* values in the chosen regions

The distribution of averaged *β* values depends on the number of individuals in each dataset and the number of CpG sites within each category. While the number of categorized CpG sites varied per each SNP (Table S1), the averaged *β* gave distributions that are all normal-shape-like (Figure 1 in the main text and Figure S3, below). The ranges of these averaged *β* values are quite narrow, but this may be attributed to each measurement being an average value of the actual methylation levels from several cells in the sample. The underlying *β* values can be directly translated to the biological effect: *β* = 0 means there is no methylation in this CpG site, while *β* =1 means this CpG site is fully methylated in all samples.

**Table S1:**
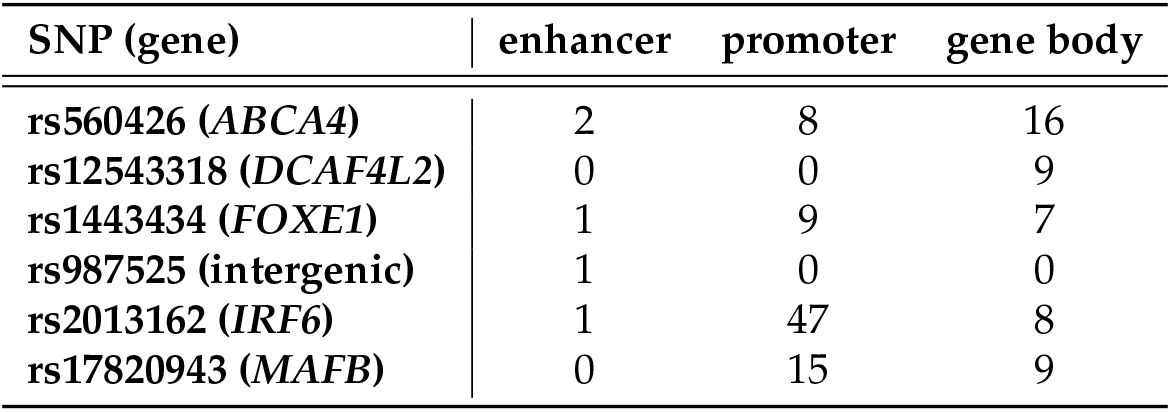
Number of CpG sites within 50 kbp from each SNP.

**Figure S3:**
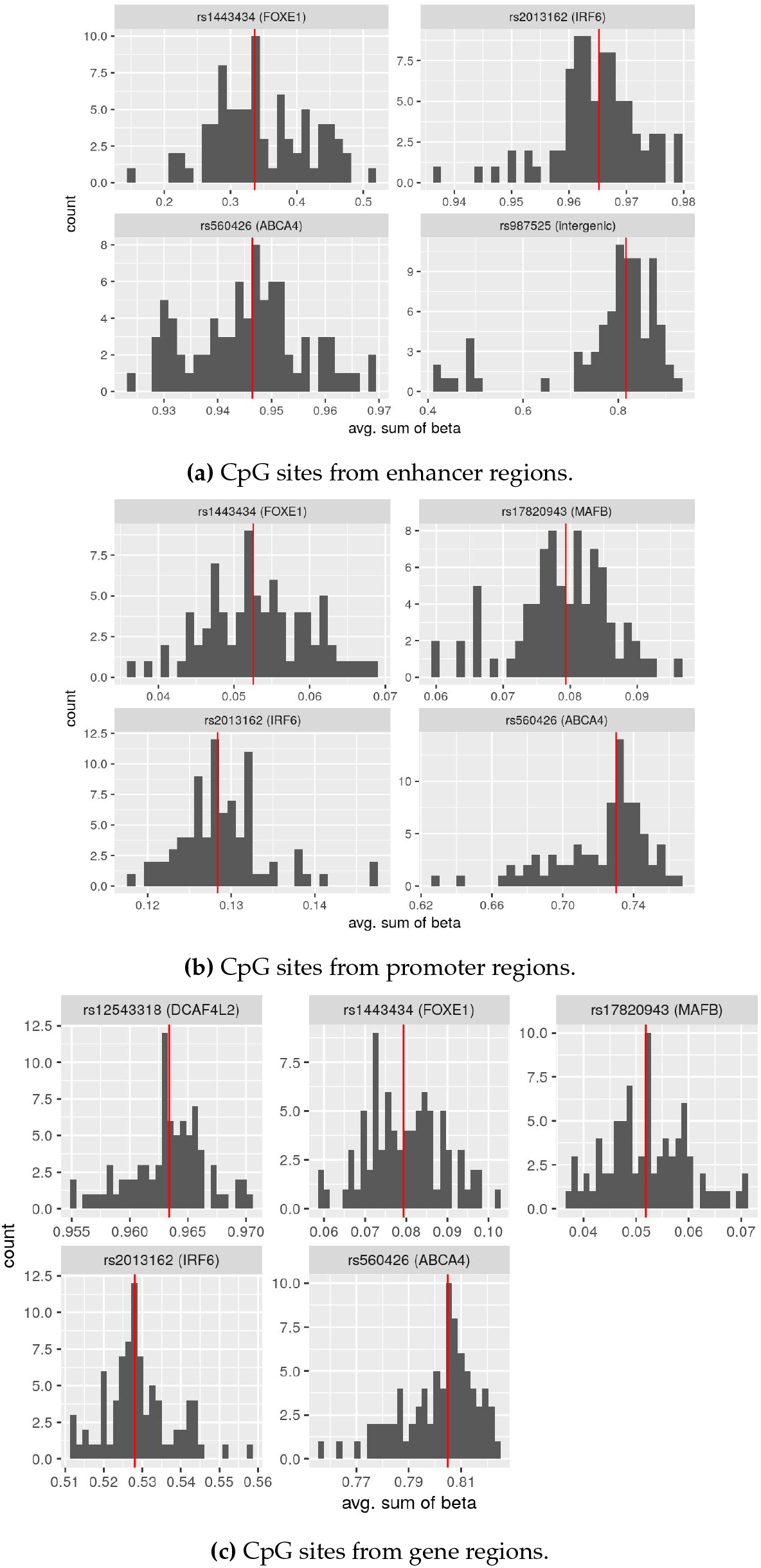
Histograms for the averaged *β* values, CLO dataset.

### S2 Main results

Below, we give all the results of the analyses: *p*-values for the G×Me or PoO×Me and trend tests, as well as several figures with the relative risks presented for specific SNPs.

#### S2.1 G×Me

##### S2.1.1 CL/P dataset

**Table S2:** The *p*-values for the G×Me and trend tests, based on the analysis of the CL/P dataset.

| SNP (gene) | PROMOTER |  | ENHANCER |  | GENE |  |
| --- | --- | --- | --- | --- | --- | --- |
| | $G \times Me$ | trend | $G \times Me$ | trend | $G \times Me$ | trend |
| rs12543318 (*DCAF4L2*) | *n.a.* * | *n.a.* | *n.a.* | *n.a.* | 0.24 | 0.31 |
| rs17820943 (*MAFB*) | 0.59 | 0.54 | *n.a.* | *n.a.* | 0.27 | 0.6 |
| rs2013162 (*IRF6*) | 0.095 | 0.03 | 0.68 | 0.49 | $2.1 \cdot 10^{-4}$ | $1.1 \cdot 10^{-4}$ |
| rs560426 (*ABCA4*) | 0.013 | 0.004 | 0.2 | 0.5 | 0.0089 | 0.0021 |
| rs987525 (intergenic) | *n.a.* | *n.a.* | $1.2 \cdot 10^{-4}$ | $3.5 \cdot 10^{-5}$ | *n.a.* | *n.a.* |

\*
itshapen.a. = not applicable (there were no CpGs within this region around this SNP)

**Figure S4:**
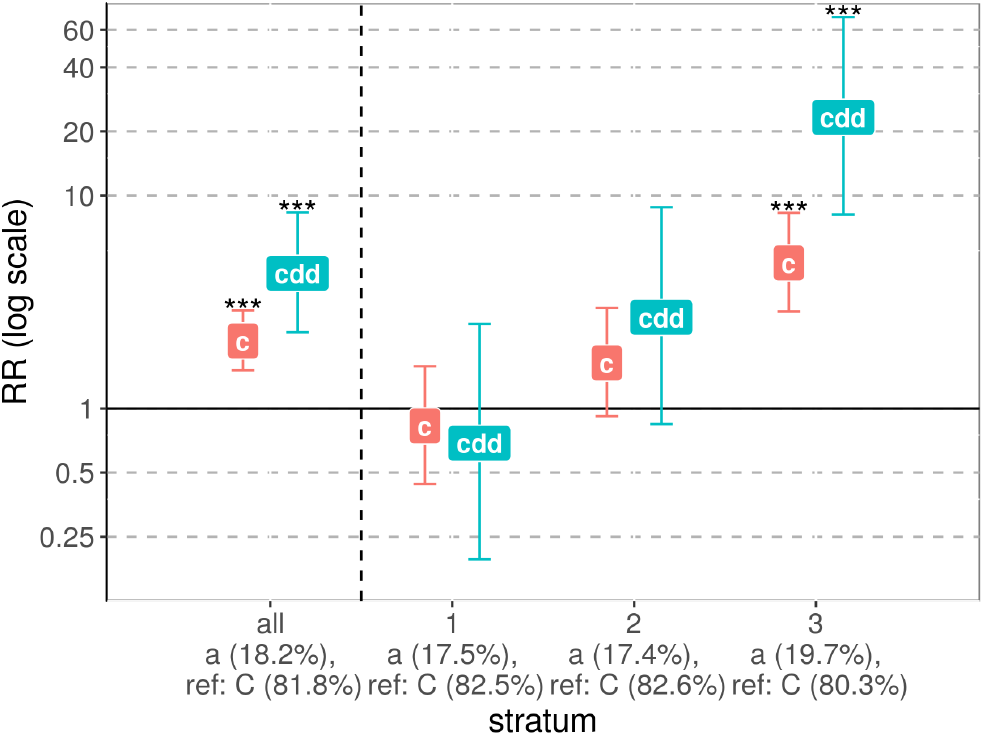
Interaction between the SNP rs987525 and methylation (G×Me) of the CpGs from the enhancer region around it. The x-axis groups the results into “all” (without stratification of the dataset) and the results for each stratum: “1” denoting low methylation level, “2” — medium, and “3” — high methylation level. The y-axis shows the relative risk (*RR*) on a log scale, with “c” denoting the child effect when only one minor allele is inherited, and “cdd” denoting the child effect of a double-dose allele inheritance.

##### S2.1.2 CLO dataset

**Table S3:** The *p*-values for the G×Me and trend tests, based on the analysis of the CLO dataset.

| SNP (gene) | PROMOTER |  | ENHANCER |  | GENE |  |
| --- | --- | --- | --- | --- | --- | --- |
|  | G×Me | trend | G×Me | trend | G×Me | trend |
| rs12543318 (*DCAF4L2*) | *n.a.* * | *n.a.* | *n.a.* | *n.a.* | 0.7 | 0.52 |
| rs1443434 (*FOXE1*) | 0.22 | 0.099 | 0.078 | 0.41 | 0.82 | 0.59 |
| rs17820943 (*MAFB*) | 0.99 | 0.92 | *n.a.* | *n.a.* | 0.88 | 0.88 |
| rs2013162 (*IRF6*) | 0.35 | 0.16 | 0.39 | 0.66 | 0.34 | 0.17 |
| rs560426 (*ABCA4*) | 0.0016 | 0.0011 | 0.66 | 0.45 | 0.0018 | $4.6 \cdot 10^{-4}$ |
| rs987525 (intergenic) | *n.a.* | *n.a.* | 0.83 | 0.68 | *n.a.* | *n.a.* |

\*
itshapen.a. = not applicable (there were no CpGs within this region around this SNP)

##### S2.1.3 Control dataset

**Table S4:** The *p*-values for the G×Me and trend tests, based on the analysis of the control dataset.

| SNP (gene) | PROMOTER |  | ENHANCER |  | GENE |  |
| --- | --- | --- | --- | --- | --- | --- |
|  | G×Me | trend | G×Me | trend | G×Me | trend |
| rs2013162 (*IRF6*) | 0.58 | 0.35 | 0.27 | 0.13 | $8.4 \cdot 10^{-5}$ | $1.6 \cdot 10^{-5}$ |
| rs560426 (*ABCA4*) | 0.67 | 0.43 | 0.012 | 0.66 | 0.92 | 0.76 |
| rs987525 (intergenic) | *n.a.* * | *n.a.* | 0.048 | 0.019 | *n.a.* | *n.a.* |

\*
itshapen.a. = not applicable (there were no CpGs within this region around this SNP)

**Figure S5:**
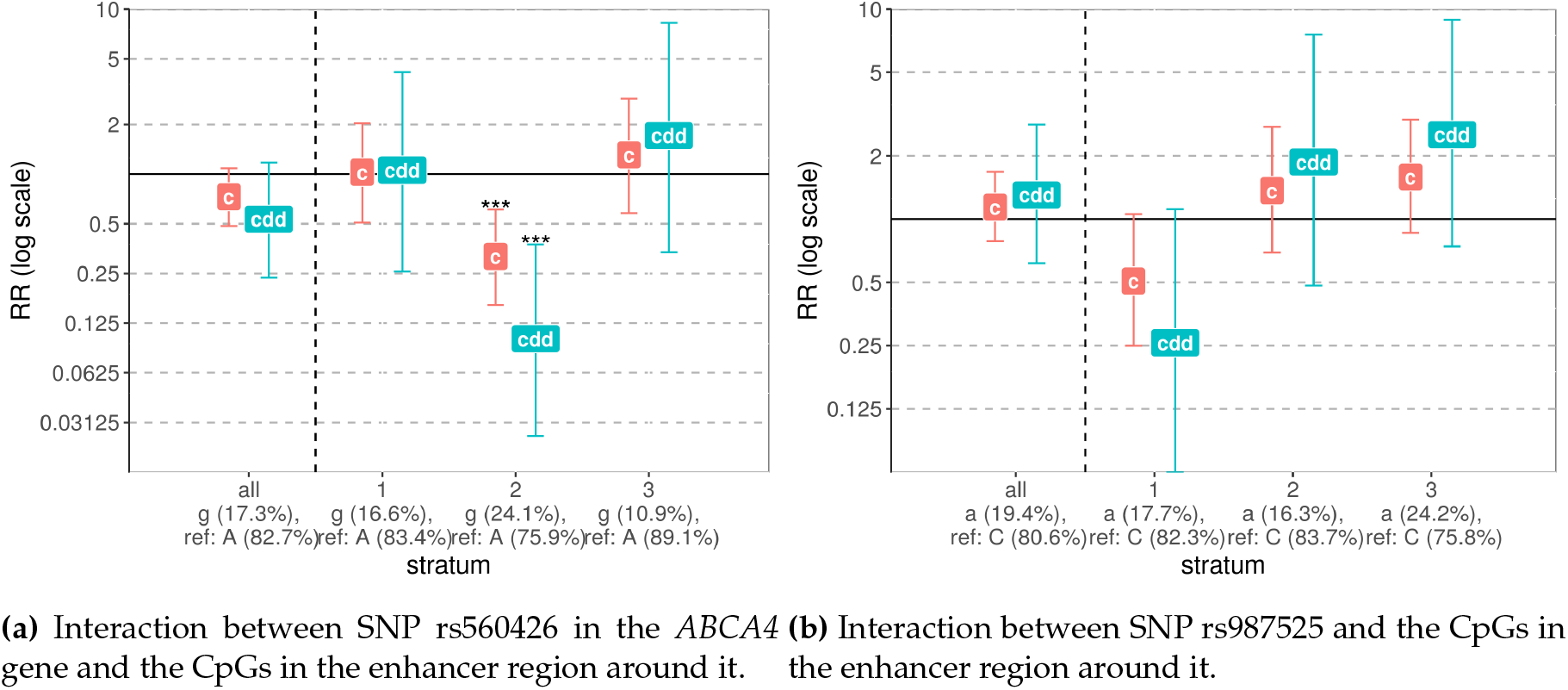
Significant G×Me interactions found in the analysis of the control dataset. The x-axis groups the results into “all” (without stratification of the dataset) and the results for each stratum: “1” denoting low methylation level, “2” — medium, and “3” — high methylation level. The y-axis shows the relative risk (*RR*) on a log scale, with “c” denoting the child effect when only one minor allele is inherited, and “cdd” denoting the child effect of a double-dose allele inheritance.

##### S2.1.4 Randomly chosen SNPs, CL/P dataset

**Table S5:** The randomly chosen SNPs and their chromosomal positions.PoO×Me

| SNP NAME | POSITION |
| --- | --- |
| rs6673129 | chr1:2202774 |
| rs9435429 | chr1:107526857 |
| rs4656971 | chr1:160921230 |
| rs10495742 | chr2:23550244 |
| rs12373638 | chr2:66731242 |
| rs12998926 | chr2:114546660 |
| rs13427642 | chr2:213562066 |
| rs6842830 | chr4:55873926 |
| rs11736543 | chr4:139696731 |
| rs9349651 | chr6:52795489 |
| rs420716 | chr6:97146770 |
| rs561690 | chr8:27510463 |
| rs7873986 | chr9:104338305 |
| rs1394371 | chr15:78724469 |
| rs4785434 | chr16:50587976 |
| rs12918462 | chr16:55127191 |
| rs8106239 | chr19:36085358 |
| rs410852 | chr19:54800371 |
| rs159099 | chr20:53631117 |
| rs2142239 | chr21:15765015 |

#### S2.2 PoO×Me

##### S2.2.1 CL/P dataset

**Table S6:** Results of the analysis of the CL/P dataset: the *p*-values for the PoO×Me interaction and trend tests.

|  | PROMOTER |  | ENHANCER |  | GENE |  |
| --- | --- | --- | --- | --- | --- | --- |
| SNP (gene) | PoO×Me | trend | PoO×Me | trend | PoO×Me | trend |
| rs12543318 (*DCAF4L2*) | *n.a.* * | *n.a.* | *n.a.* | *n.a.* | 0.025 | 0.011 |
| rs17820943 (*MAFB*) | 0.95 | 0.83 | *n.a.* | *n.a.* | 0.17 | 0.14 |
| rs2013162 (*IRF6*) | 0.48 | 0.53 | 0.83 | 0.66 | 0.99 | 0.92 |
| rs560426 (*ABCA4*) | 0.99 | 0.95 | 0.2 | 0.088 | 0.23 | 0.8 |
| rs987525 (intergenic) | *n.a.* | *n.a.* | 0.38 | 0.35 | *n.a.* | *n.a.* |

\* *n.a.* = not applicable (there were no CpGs within this region around this SNP)

##### S2.2.2 CLO dataset

**Table S7:** Results of the analysis of the CLO dataset: the *p*-values for the PoO×Me interaction and trend tests.

|  | PROMOTER |  | ENHANCER |  | GENE |  |
| --- | --- | --- | --- | --- | --- | --- |
| SNP (gene) | PoO×Me | trend | PoO×Me | trend | PoO×Me | trend |
| rs12543318 (*DCAF4L2*) | *n.a.* * | *n.a.* | *n.a.* | *n.a.* | 0.4 | 0.31 |
| rs1443434 (*FOXE1*) | 0.74 | 0.84 | 0.51 | 0.6 | 0.75 | 0.82 |
| rs17820943 (*MAFB*) | 0.71 | 0.47 | *n.a.* | *n.a.* | 0.24 | 0.14 |
| rs2013162 (*IRF6*) | 0.5 | 0.93 | 0.37 | 0.36 | 0.61 | 0.85 |
| rs560426 (*ABCA4*) | 0.61 | 0.33 | 0.68 | 0.38 | 0.32 | 0.15 |
| rs987525 (intergenic) | *n.a.* | *n.a.* | 0.69 | 0.42 | *n.a.* | *n.a.* |

\* *n.a.* = not applicable (there were no CpGs within this region around this SNP)

##### S2.2.3 Control dataset

**Table S8:** Results of the analysis of the control dataset: the *p*-values for the PoO×Me interaction and trend tests.

|  | PROMOTER |  | ENHANCER |  | GENE |  |
| --- | --- | --- | --- | --- | --- | --- |
| SNP (gene) | PoO×Me | trend | PoO×Me | trend | PoO×Me | trend |
| rs2013162 (*IRF6*) | 0.73 | 0.68 | 0.066 | 0.57 | 0.77 | 0.92 |
| rs560426 (*ABCA4*) | 0.49 | 0.39 | 0.95 | 0.74 | 0.42 | 0.24 |
| rs987525 (intergenic) | *n.a.* * | *n.a.* | 0.08 | 0.98 | *n.a.* | *n.a.* |

\* *n.a.* = not applicable (there were no CpGs within this region around this SNP)

##### S2.2.4 PoO scan, CL/P dataset

**Table S9.**
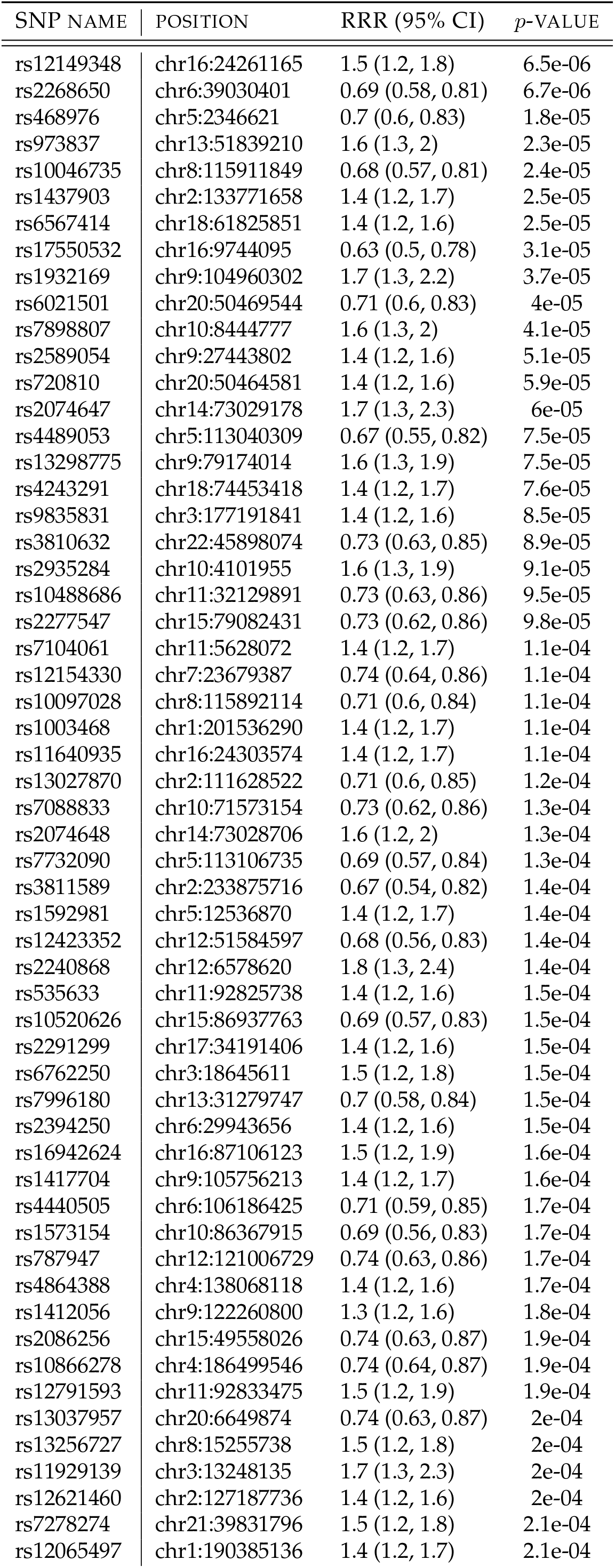
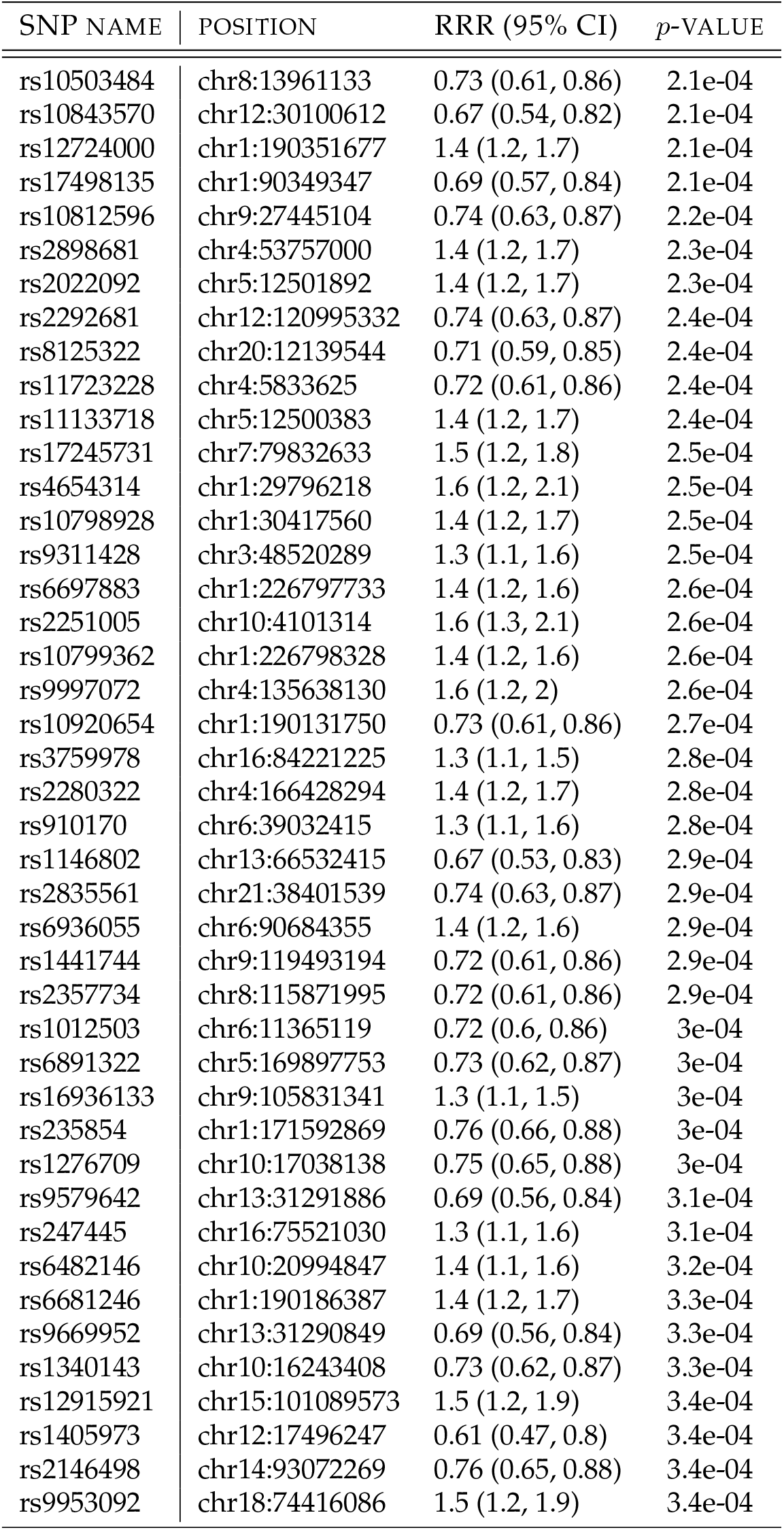
Top 100 SNPs from the PoO scan of the genotype CL/P dataset.

| SNP NAME | POSITION | RRR (95% CI) | p-VALUE |
| --- | --- | --- | --- |
| rs12149348 | chr16:24261165 | 1.5 (1.2, 1.8) | 6.5e-06 |
| rs2268650 | chr6:39030401 | 0.69 (0.58, 0.81) | 6.7e-06 |
| rs468976 | chr5:2346621 | 0.7 (0.6, 0.83) | 1.8e-05 |
| rs973837 | chr13:51839210 | 1.6 (1.3, 2) | 2.3e-05 |
| rs10046735 | chr8:115911849 | 0.68 (0.57, 0.81) | 2.4e-05 |
| rs1437903 | chr2:133771658 | 1.4 (1.2, 1.7) | 2.5e-05 |
| rs6567414 | chr18:61825851 | 1.4 (1.2, 1.6) | 2.5e-05 |
| rs17550532 | chr16:9744095 | 0.63 (0.5, 0.78) | 3.1e-05 |
| rs1932169 | chr9:104960302 | 1.7 (1.3, 2.2) | 3.7e-05 |
| rs6021501 | chr20:50469544 | 0.71 (0.6, 0.83) | 4e-05 |
| rs7898807 | chr10:8444777 | 1.6 (1.3, 2) | 4.1e-05 |
| rs2589054 | chr9:27443802 | 1.4 (1.2, 1.6) | 5.1e-05 |
| rs720810 | chr20:50464581 | 1.4 (1.2, 1.6) | 5.9e-05 |
| rs2074647 | chr14:73029178 | 1.7 (1.3, 2.3) | 6e-05 |
| rs4489053 | chr5:113040309 | 0.67 (0.55, 0.82) | 7.5e-05 |
| rs13298775 | chr9:79174014 | 1.6 (1.3, 1.9) | 7.5e-05 |
| rs4243291 | chr18:74453418 | 1.4 (1.2, 1.7) | 7.6e-05 |
| rs9835831 | chr3:177191841 | 1.4 (1.2, 1.6) | 8.5e-05 |
| rs3810632 | chr22:45898074 | 0.73 (0.63, 0.85) | 8.9e-05 |
| rs2935284 | chr10:4101955 | 1.6 (1.3, 1.9) | 9.1e-05 |
| rs10488686 | chr11:32129891 | 0.73 (0.63, 0.86) | 9.5e-05 |
| rs2277547 | chr15:79082431 | 0.73 (0.62, 0.86) | 9.8e-05 |
| rs7104061 | chr11:5628072 | 1.4 (1.2, 1.7) | 1.1e-04 |
| rs12154330 | chr7:23679387 | 0.74 (0.64, 0.86) | 1.1e-04 |
| rs10097028 | chr8:115892114 | 0.71 (0.6, 0.84) | 1.1e-04 |
| rs1003468 | chr1:201536290 | 1.4 (1.2, 1.7) | 1.1e-04 |
| rs11640935 | chr16:24303574 | 1.4 (1.2, 1.7) | 1.1e-04 |
| rs13027870 | chr2:111628522 | 0.71 (0.6, 0.85) | 1.2e-04 |
| rs7088833 | chr10:71573154 | 0.73 (0.62, 0.86) | 1.3e-04 |
| rs2074648 | chr14:73028706 | 1.6 (1.2, 2) | 1.3e-04 |
| rs7732090 | chr5:113106735 | 0.69 (0.57, 0.84) | 1.3e-04 |
| rs3811589 | chr2:233875716 | 0.67 (0.54, 0.82) | 1.4e-04 |
| rs1592981 | chr5:12536870 | 1.4 (1.2, 1.7) | 1.4e-04 |
| rs12423352 | chr12:51584597 | 0.68 (0.56, 0.83) | 1.4e-04 |
| rs2240868 | chr12:6578620 | 1.8 (1.3, 2.4) | 1.4e-04 |
| rs535633 | chr11:92825738 | 1.4 (1.2, 1.6) | 1.5e-04 |
| rs10520626 | chr15:86937763 | 0.69 (0.57, 0.83) | 1.5e-04 |
| rs2291299 | chr17:34191406 | 1.4 (1.2, 1.6) | 1.5e-04 |
| rs6762250 | chr3:18645611 | 1.5 (1.2, 1.8) | 1.5e-04 |
| rs7996180 | chr13:31279747 | 0.7 (0.58, 0.84) | 1.5e-04 |
| rs2394250 | chr6:29943656 | 1.4 (1.2, 1.6) | 1.5e-04 |
| rs16942624 | chr16:87106123 | 1.5 (1.2, 1.9) | 1.6e-04 |
| rs1417704 | chr9:105756213 | 1.4 (1.2, 1.7) | 1.6e-04 |
| rs4440505 | chr6:106186425 | 0.71 (0.59, 0.85) | 1.7e-04 |
| rs1573154 | chr10:86367915 | 0.69 (0.56, 0.83) | 1.7e-04 |
| rs787947 | chr12:121006729 | 0.74 (0.63, 0.86) | 1.7e-04 |
| rs4864388 | chr4:138068118 | 1.4 (1.2, 1.6) | 1.7e-04 |
| rs1412056 | chr9:122260800 | 1.3 (1.2, 1.6) | 1.8e-04 |
| rs2086256 | chr15:49558026 | 0.74 (0.63, 0.87) | 1.9e-04 |
| rs10866278 | chr4:186499546 | 0.74 (0.64, 0.87) | 1.9e-04 |
| rs12791593 | chr11:92833475 | 1.5 (1.2, 1.9) | 1.9e-04 |
| rs13037957 | chr20:6649874 | 0.74 (0.63, 0.87) | 2e-04 |
| rs13256727 | chr8:15255738 | 1.5 (1.2, 1.8) | 2e-04 |
| rs11929139 | chr3:13248135 | 1.7 (1.3, 2.3) | 2e-04 |
| rs12621460 | chr2:127187736 | 1.4 (1.2, 1.6) | 2e-04 |
| rs7278274 | chr21:39831796 | 1.5 (1.2, 1.8) | 2.1e-04 |
| rs12065497 | chr1:190385136 | 1.4 (1.2, 1.7) | 2.1e-04 |
| rs10503484 | chr8:13961133 | 0.73 (0.61, 0.86) | 2.1e-04 |
| rs10843570 | chr12:30100612 | 0.67 (0.54, 0.82) | 2.1e-04 |
| rs12724000 | chr1:190351677 | 1.4 (1.2, 1.7) | 2.1e-04 |
| rs17498135 | chr1:90349347 | 0.69 (0.57, 0.84) | 2.1e-04 |
| rs10812596 | chr9:27445104 | 0.74 (0.63, 0.87) | 2.2e-04 |
| rs2898681 | chr4:53757000 | 1.4 (1.2, 1.7) | 2.3e-04 |
| rs2022092 | chr5:12501892 | 1.4 (1.2, 1.7) | 2.3e-04 |
| rs2292681 | chr12:120995332 | 0.74 (0.63, 0.87) | 2.4e-04 |
| rs8125322 | chr20:12139544 | 0.71 (0.59, 0.85) | 2.4e-04 |
| rs11723228 | chr4:5833625 | 0.72 (0.61, 0.86) | 2.4e-04 |
| rs11133718 | chr5:12500383 | 1.4 (1.2, 1.7) | 2.4e-04 |
| rs17245731 | chr7:79832633 | 1.5 (1.2, 1.8) | 2.5e-04 |
| rs4654314 | chr1:29796218 | 1.6 (1.2, 2.1) | 2.5e-04 |
| rs10798928 | chr1:30417560 | 1.4 (1.2, 1.7) | 2.5e-04 |
| rs9311428 | chr3:48520289 | 1.3 (1.1, 1.6) | 2.5e-04 |
| rs6697883 | chr1:226797733 | 1.4 (1.2, 1.6) | 2.6e-04 |
| rs2251005 | chr10:4101314 | 1.6 (1.3, 2.1) | 2.6e-04 |
| rs10799362 | chr1:226798328 | 1.4 (1.2, 1.6) | 2.6e-04 |
| rs9997072 | chr4:135638130 | 1.6 (1.2, 2) | 2.6e-04 |
| rs10920654 | chr1:190131750 | 0.73 (0.61, 0.86) | 2.7e-04 |
| rs3759978 | chr16:84221225 | 1.3 (1.1, 1.5) | 2.8e-04 |
| rs2280322 | chr4:166428294 | 1.4 (1.2, 1.7) | 2.8e-04 |
| rs910170 | chr6:39032415 | 1.3 (1.1, 1.6) | 2.8e-04 |
| rs1146802 | chr13:66532415 | 0.67 (0.53, 0.83) | 2.9e-04 |
| rs2835561 | chr21:38401539 | 0.74 (0.63, 0.87) | 2.9e-04 |
| rs6936055 | chr6:90684355 | 1.4 (1.2, 1.6) | 2.9e-04 |
| rs1441744 | chr9:119493194 | 0.72 (0.61, 0.86) | 2.9e-04 |
| rs2357734 | chr8:115871995 | 0.72 (0.61, 0.86) | 2.9e-04 |
| rs1012503 | chr6:11365119 | 0.72 (0.6, 0.86) | 3e-04 |
| rs6891322 | chr5:169897753 | 0.73 (0.62, 0.87) | 3e-04 |
| rs16936133 | chr9:105831341 | 1.3 (1.1, 1.5) | 3e-04 |
| rs235854 | chr1:171592869 | 0.76 (0.66, 0.88) | 3e-04 |
| rs1276709 | chr10:17038138 | 0.75 (0.65, 0.88) | 3e-04 |
| rs9579642 | chr13:31291886 | 0.69 (0.56, 0.84) | 3.1e-04 |
| rs247445 | chr16:75521030 | 1.3 (1.1, 1.6) | 3.1e-04 |
| rs6482146 | chr10:20994847 | 1.4 (1.1, 1.6) | 3.2e-04 |
| rs6681246 | chr1:190186387 | 1.4 (1.2, 1.7) | 3.3e-04 |
| rs9669952 | chr13:31290849 | 0.69 (0.56, 0.84) | 3.3e-04 |
| rs1340143 | chr10:16243408 | 0.73 (0.62, 0.87) | 3.3e-04 |
| rs12915921 | chr15:101089573 | 1.5 (1.2, 1.9) | 3.4e-04 |
| rs1405973 | chr12:17496247 | 0.61 (0.47, 0.8) | 3.4e-04 |
| rs2146498 | chr14:93072269 | 0.76 (0.65, 0.88) | 3.4e-04 |
| rs9953092 | chr18:74416086 | 1.5 (1.2, 1.9) | 3.4e-04 |

### S3 G × Me analyses using four strata

**Figure S6:**
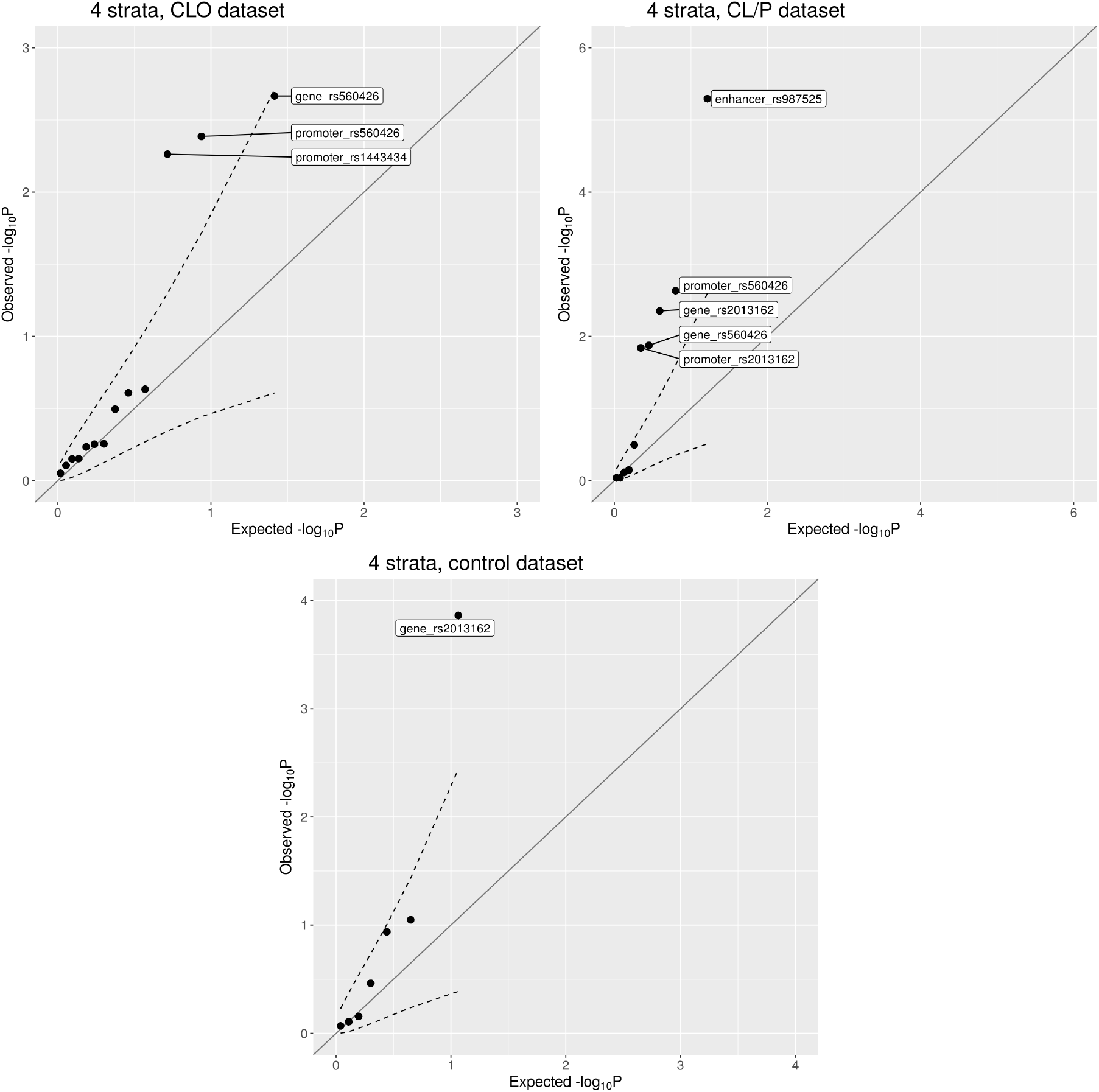
Quantile-quantile plots of the interaction *p*-values from the G×Me analyses, using four equally sized strata; the dashed lines represent the 95% confidence interval.

